# A resource for exploring the understudied human kinome for research and therapeutic opportunities

**DOI:** 10.1101/2020.04.02.022277

**Authors:** Nienke Moret, Changchang Liu, Benjamin M. Gyori, John A. Bachman, Albert Steppi, Clemens Hug, Rahil Taujale, Liang-Chin Huang, Matthew E. Berginski, Shawn M. Gomez, Natarajan Kannan, Peter K. Sorger

**Affiliations:** The NIH Understudied Kinome Consortium; Laboratory of Systems Pharmacology, Department of Systems Biology, Harvard Program in Therapeutic Science, Harvard Medical School, Boston, Massachusetts 02115, USA; Institute of Bioinformatics, University of Georgia, Athens, GA, 30602 USA; Department of Pharmacology, The University of North Carolina at Chapel Hill, Chapel Hill, NC 27599, USA; Joint Department of Biomedical Engineering at the University of North Carolina at Chapel Hill and North Carolina State University, Chapel Hill, NC 27599, USA

**Keywords:** cancer, cheminformatics, drug discovery, human kinome, kinase inhibitors

## Abstract

The functions of protein kinases have been widely studied and over 60 kinase inhibitors are FDA-approved drugs. Membership in the human kinome is nonetheless subject to multiple overlapping and inconsistent definitions and is unevenly studied, complicating functional genomics and chemical genetics. We describe objective criteria for refining the definition of the human kinome to comprise an extended set of 710 kinase domains and a more narrowly curated set of 557 protein kinase like (PKL) domains. An online tool (www.kinome.org) makes it possible to sort these sets on multiple structural and functional criteria. Focusing on the least studied one-third of the kinome we find that many proteins are differentially expressed, essential in multiple cell lines, and mutated in the Cancer Genome Atlas. We show that some understudied kinases are high affinity off-targets of clinical-grade compounds and approved drugs and we describe an optimized small molecule library making use of this information for selective kinome perturbation. We conclude that the understudied kinome contains physiologically important proteins, including possible targets for future drug discovery campaigns.

## INTRODUCTION

*Note – This manuscript includes three text boxes that are found after the results section*

Protein phosphorylation is widespread in eukaryotic cells (Cohen, 2002) and mediates many critical events in cell fate determination, cell cycle control and signal transduction (Hunter, 1995). The 3D structures (protein folds) (Knighton et al., 1991a) and catalytic activities (Adams, 2001) of eukaryotic protein kinases (ePKs), of which ∼500 are found in humans (Manning et al., 2002), have been intensively investigated for many years: to date, structures for over 280 unique domains and ∼4,000 co-complexes have been deposited in the PDB database. The fold of ePKs is thought to have arisen in prokaryotes (Lai et al., 2016) and evolved to include tyrosine kinases in metazoans (Darnell, 1997; Wijk and Snel, 2020), resulting in a diverse set of enzymes (Hanks and Hunter, 1995; Hanks et al., 1988) that are often linked—in a single protein—to other functional protein domains such as SH2 and SH3 peptide-binding motifs. In addition, 13 human proteins have two ePK kinase domains. A recent review describes the structural properties of ePKs and the drugs that bind them (Kanev et al., 2019).

With an accessible binding pocket and demonstrated involvement in many disease pathways, protein kinases are attractive drug targets (Bhullar et al., 2018). Protein kinase inhibitors, and the few activators that have been identified (e.g., AMPK activation by salicylate and A-769662 (Hawley et al., 2012)), are diverse in structure and mechanism. These molecules include a large family of ATP-competitive inhibitors that bind in the enzyme active site and a smaller number of non-competitive “allosteric” inhibitors that bind elsewhere as well as small molecule PROTAC degraders whose binding to a kinase promotes ubiquitin-dependent degradation (Jones, 2018). Multiple antibodies that target the ligand binding sites of receptor kinases or that interfere with a receptor’s ability to homo or hetero-oligomerize have been developed into human therapeutics (FAUVEL and Yasri, 2014). Small molecule kinase inhibitors have been intensively studied in human clinical trials and over 60 have been developed into FDA-approved drugs (Kanev et al., 2019).

Despite the general importance of kinases in cellular physiology as well as their druggability and frequent mutation in disease, a substantial subset of the kinome has been relatively overlooked with respect to the number of publications and public research funding. This has given rise to a project within the NIH’s *Illuminating the Druggable Genome* Program (IDG) (Finan et al., 2017), to investigate the relatively understudied “dark kinome” and identify proteins with roles in human biology and disease. The NIH has distributed a preliminary list of understudied kinases based on the number of discoverable publications and the presence/absence of grant (NIH R01) funding and we and others are developing reagents to study the properties of these enzymes (Berginski et al., 2020; Huang et al., 2018). Defining which kinases are understudied and their relationship to other, better characterized kinases, necessarily involves a working definition of the full kinome and a survey of the current state of knowledge.

Establishing the membership of the human kinome is a more subtle task than might be expected. The first human “kinome” was defined as the family of proteins homologous to enzymes shown experimentally to have peptide-directed phosphotransferase activity. A groundbreaking 2002 paper by Manning et al. (Manning et al., 2002) defined a kinome having 514 members. This list has subsequently been updated via the <u>KinHub</u> Web resource (Eid et al., 2017a) to include 522 proteins. The kinase domain of protein Kinase A (PKA), a hetero-oligomer of a regulatory and a catalytic subunit, was the first to be crystallized and is often regarded as the prototype of the ePK fold (Knighton et al., 1991a, 1991b). This fold involves two distinct lobes with an ATP-binding catalytic cleft lying between the lobes that is characterized, at the level of primary sequence, by 12 recurrent elements with a total of ∼30 highly conserved residues (Lai et al., 2016). Of 514 proteins in the human kinome as defined by Manning et al, 478 have an ePK fold.

Kinases have diverged in multiple ways to generate protein subfamilies distinct in sequence and structure from PKA including kinases with so-called eukaryotic like folds, atypical folds and unrelated folds. The eukaryotic like kinases (eLKs) are similar to ePKs in that they retain significant sequence similarity to the N-terminal region of ePKs but differ in the substrate binding lobe. TP53RK, a regulator of p53 (TP53) (Abe et al., 2001) is an example of a serine/threonine protein kinase with an eLK fold. Kinases with an atypical fold (aPKs) are distinct from ePKs and eLKs in that they have weak sequence similarity to ePKs but nevertheless adopt an ePK-like three-dimensional structure. aPKs include some well-studied protein kinases such as the DNA damage sensing ATM and ATR kinases (Abraham, 2001). The aPK and eLK folds are not limited to protein kinases. The family of PI3K lipid kinases, some of the most heavily mutated genes in breast cancer (Mukohara, 2015), also adopt an aPK fold (Kanev et al., 2019). Similarly, the choline kinase CHKA, a key player in dysregulated choline metabolism in cancer cells and a chemotherapy target, adopts the eLK fold (Glunde et al., 2011; Oruganty et al., 2016). Over 200 additional proteins are annotated as “kinase” in UniProt. Among these proteins, 141 have structures unrelated to the protein kinase fold and they are therefore termed uPKs (unrelated to Protein Kinases). Enzymes with phosphotransferase activity in the uPK family include hexokinases that phosphorylate sugars, but also proteins with bromodomains (e.g., BRD2, BRD3 and BRD4). Multiple uPK proteins, including those with bromodomains, bind to small molecule kinase inhibitors (Ciceri et al., 2014) making it useful, from the perspective of drug discovery, to study kinase-like proteins at the same time as kinases themselves.

While protein kinases could in principle be defined strictly as enzymes that catalyze phospho-transfer from ATP onto serine, threonine and tyrosine, such a definition would exclude structurally and functionally related lipid kinases as well as many of the protein families relevant to drug discovery. It would also fail to account for a lack of functional data for a substantial number of proteins, potentially excluding from consideration kinases that are physiologically or catalytically active but have not yet been tested in biochemical assays. This problem of missing biochemical data has resulted in kinome definitions that rely on sequence alignment and structural data to identify closely related folds (Ciceri et al., 2014); in these definitions, uPKs having kinase activity as well as bromodomains that potently bind and are inactivated by kinase inhibitors are often overlooked. Moreover, as Hidden Markov Models (HMMs) and other ways of recognizing kinase homology have become more sophisticated, additional proteins have been added to the kinome (Briedis et al., 2008).

Kinases that phosphorylate molecules other than proteins or peptides, such as lipids or metabolites, are frequently components of signal transduction pathways involving kinase cascades (Mosca et al., 2012). Phosphoinositide 3-kinases that modify lipids with a role as secondary messengers are a well-known example (Verheijen et al., 2011). Moreover, some metabolic kinases have been shown to be active against peptide or protein substrates (Lu and Hunter, 2018), challenging the conventional notion that small molecule and peptide-directed kinases are distinct families of enzymes. One example is the pyruvate kinase PKM2, which in addition to its well-known function in generating pyruvate from phosphoenolpyruvate, can also phosphorylate histone H3 at T11, thereby activating transcription downstream of EGFR-signaling (Yang et al., 2012). Considered together, these findings suggest that it is important to define the kinome along multiple axes, based on fold or sequence homology, ability to bind small molecule kinase inhibitors, and biochemical function.

The published literature uses multiple different definitions of the human kinome, likely because there is no consensus on which kinase set is superior. The functional definition of the kinome can often also differ depending on the research application. “Omics” studies and computational studies benefit from an expansive definition of the kinome encompassing all related enzymes. For example, kinome-wide profiling typically involves screening compounds against panels of recombinant enzymes (e.g., KINOMEscan (Posy et al., 2011)) or chemo-proteomics in which competitive binding to ATP-like ligands on beads (so-called kinobeads (Klaeger et al., 2017) or multiplexed inhibitor beads - MIBs (Cousins et al., 2018)) are assayed using mass spectrometry. In contrast, screens for kinases that phosphorylate a specific protein sequence, for example, would logically focus on enzymes that are known, or are likely based on structure, to have peptide-directed phosphotransferase activity.

In this paper we use data-driven criteria to generate new sets of kinases having different inclusion and exclusion criteria. We quantify the mechanistic knowledge available for each kinase domain using machine reading and automatic network assembly with the INDRA software (Gyori et al., 2017) and compute membership in the understudied kinome, based on current mechanistic knowledge. We then consolidate available data on kinase activity and function with the goal of determining which understudied kinases merit additional attention. Functional evidence in this context is typically indirect and includes data from TCGA (The Cancer Genome Atlas, (Weinstein et al., 2013)) on the frequency with which a kinase is mutated in a particular type of cancer. In aggregate, the evidence strongly suggests that the understudied kinome is likely to contain multiple enzymes worthy of in-depth study, a subset of which may be viable therapeutic targets. All of the information in this manuscript is available in supplementary materials and interactively through www.kinome.org. Additional information on the understudied kinome can be found on a related resource at www.darkkinome.org (Berginski et al., 2020).

## RESULTS

### The composition of the human kinome

An initial set of human kinases was obtained from Manning et al. (Manning et al., 2002) (referred to below as “Manning”) and a second from Eid et al. (Eid et al., 2017a) via the <u>KinHub</u> Web resource; a set of dark kinases according to the IDG was obtained from the NIH solicitation (updated in January 2018, referred to below as “IDG dark kinases”) and a fourth set of all 684 proteins tagged to have “kinase activity” was obtained from UniProt. The UniProt set includes protein kinases, lipid kinases and other small molecule kinases. These sets are overlapping but not identical (Figure 1A). For example, eight IDG dark kinases that are absent from Manning and KinHub (CSNK2A3, PIK3C2B, PIK3C2G, PIP4K2C, PI4KA, PIP5K1A, PIP5K1B, and PIP5K1C) are found in the UniProt list. We assembled a superset of 710 domains (the “extended kinome”) that includes all kinases from the four sources listed above and then used curated alignment profiles (HMM models) and structural analysis (Kannan et al., 2007) to subdivide domains into three primary categories: “Protein Kinase Like” (PKL), if the kinase domain was similar to known protein kinases in sequence and 3D-structure; “Unrelated to Protein Kinase” (uPK), if the kinase domain was distinct from known protein kinases; and “Unknown” if there was insufficient information to decide (see methods) (Kannan et al., 2007). PKLs were further subdivided into eukaryotic protein kinases (ePKs, discussed in the introduction), eukaryotic like kinases (eLKs) and kinases with an atypical fold (aPKs) as previously described (Kannan and Neuwald, 2005; Kannan et al., 2007). ePKs and eLKs share detectable sequence similarity in the ATP binding lobe and some portions of the substrate binding lobe (up to the conserved F-helix (Kannan et al., 2007)). aPKs, on the other hand, display no significant sequence similarity to ePKs and eLKs, but nevertheless adopt the canonical protein kinase fold in three dimensions. Most aPKs lack the canonical F-helix aspartate in the substrate binding lobe, but share structural similarities with ePKs and eLKs in the ATP binding lobe (Figure 1B). Unfortunately, the nomenclature used in making these distinctions is not consistent across sources. In this paper, aPK refers to a subset of PKLs defined by fold and sequence similarity; this is distinct from the so-called “atypical protein kinase group” (AKGs). Proteins in the AKG include protein kinases such as ATM and ATR as well as bromo-domains and TRIM proteins (see below).

**Figure 1.**
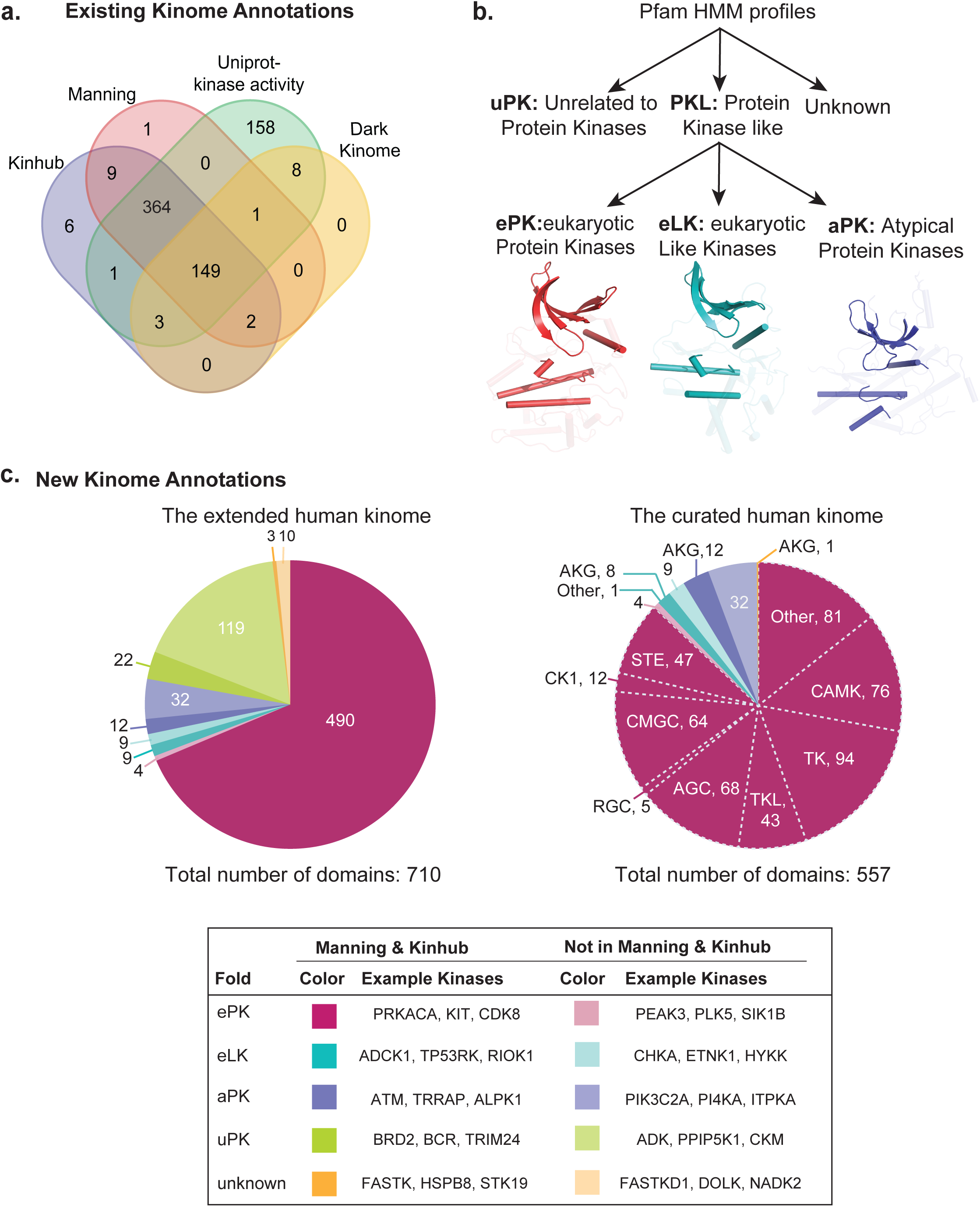
Composition of the human kinome. **(A)** Venn diagram showing the overlap in domains curated as being part of the human kinome depending on the sources. KinHub (purple) refers to a list of kinases described by Eid et al.(Eid et al., 2017b) and is accessible via http://kinhub.org/kinases.html; Manning (red) refers to the gene list prepared by Manning et al. in 2002 (Manning et al., 2002); Uniprot kinase activity (green) refers to a list of genes annotated as having kinase activity in the Uniprot database (2019) (see methods and Table S1); Dark Kinome (yellow) refers to a list of 168 understudied kinases as defined by the IDG consortium. **(B)** Schematic workflow showing how kinases are classified based on kinase three dimensional fold and sequence alignment: PKL – the *protein kinase like* fold most similar to that of the canonical PKA kinase; uPK – folds *unrelated to protein kinases* – but with significant sequence homology and known to encompass kinases against non-protein substrates as well as a limited number of protein kinases. PKLs are further subclassified into eukaryotic Protein Kinases (PKs), eukaryotic Like Kinases (eLKs) and Atypical Kinases (AKs) based on structural properties and sequence alignment. HMM refers to a Hidden Markov Model used to perform classification; see methods for details. **(C)** Pie chart showing the breakdown of 710 domains in the extended human kinome or the 557 domains in the curated kinome as defined in the current work. Subfamilies of kinases (as conventionally defined) (Manning et al., 2002) are denoted by the white dotted lines: <u>CAMK</u> – Calcium and Calmodulin regulated kinases; <u>TK</u> – Tyrosine kinase; <u>TKL</u> – Tyrosine kinase like <u>ACG</u> – named after the initials for kinases within the group PKA, PKC and PKG; CMGC – named after the initials of some members; <u>CK1</u> – cell kinase group; <u>AKG</u> – atypical protein kinase group. Legend below lists some exemplary kinases from each category. Full details can be found in **Supplementary Table S1.**

As noted previously (Garrett et al., 2011; Manning et al., 2002), structural, sequence-based and functional classifications of kinases are ambiguous and overlapping. For example, the ATM aPK is known to phosphorylate proteins DYRK2, MDM2, MDM4 and TP53 (Jassal et al., 2020) when activated by DNA double-strand breaks and it is also a member of the six-protein family of phosphatidylinositol 3-kinase-related protein kinases (PIKKs). The PIKK family has a protein fold significantly similar to lipid kinases in the PI3K/PI4K family but PI4K2A, for example, modifies phosphatidylinositol lipids and not proteins (Baumlova et al., 2014). Thus, even after use of HMM models and comparison of 3D structures, some manual curation of the kinome is necessary. To enable selection of kinases along multiple overlapping inclusion and exclusion criteria we created an interactive web-resource, available via www.kinome.org, that makes it possible to generate a wide variety of kinase sets based on user-specific criteria. This information is also available in **Supplemental Table S1**.

An extended kinome comprising 710 proteins has the disadvantage that it is substantially larger than the 525-550 domains commonly used in the literature as the set of human protein kinases. Additionally, it is frequently unknown if proteins in the extended set have phospho-transfer activity, and if so, whether it is directed against peptides, small molecules or both (Lu and Hunter, 2018). We therefore created a “curated kinome” set that closely matches the spirit of the Manning kinome definition but based on contemporary HMM models and activity data. This curated set comprises 557 kinase domains (544 genes) and includes all 556 PKLs plus STK19 (STK19 was included despite its having an “unknown fold” since it mediates serine/threonine phosphorylation of peptide substrates (Yin et al., 2019)) (**Supplemental Table S2)**. The curated kinome omits 15 uPKs found in Manning and 22 found in KinHub (including multiple TRIM family proteins (Reymond et al., 2001)) that regulate and are regulated by kinases (Ozato et al., 2008), but have no known intrinsic kinase activity; bromodomains are also omitted. The curated kinome is compared to the Manning set in Figure 1C and **Figure S1A.**

The traditional kinome tree as used by Coral (Metz et al., 2018) is not optimized to visualize the curated kinome since it primarily focused on kinases having an ePK fold. The omission of two groups of kinases is particularly problematic: i) kinases that are not in the Manning set but do have an ePK fold and were simply overlooked and ii) AKG kinases, which are often depicted as a list of proteins unrelated to each other (Metz et al., 2018). We therefore generated a new depiction of the kinome based on HMMs (see methods) that addresses these issues (Figure 2). The new depiction has two separate kinome trees – a primary tree slightly modified from the existing Coral tree (Metz et al., 2018) and a second, newly drawn tree. The primary tree was initially generated based on sequence-comparisons performed by Manning et al in 2002 and then modified to create an aesthetically pleasing representation that can be accessed via the Coral web application (Metz et al., 2018). The organization of the Coral tree (including the roots for various ePKs subfamilies) is supported by multiple lines of structural, sequence and phylogenetic evidence and generally regarded as an objective representation of the evolutionary relationships of different kinase subfamilies. We have updated the tree to include four additional kinases with an ePK fold (CSNK2A3, PEAK3, PLK5, SIK1B).

**Figure 2.**
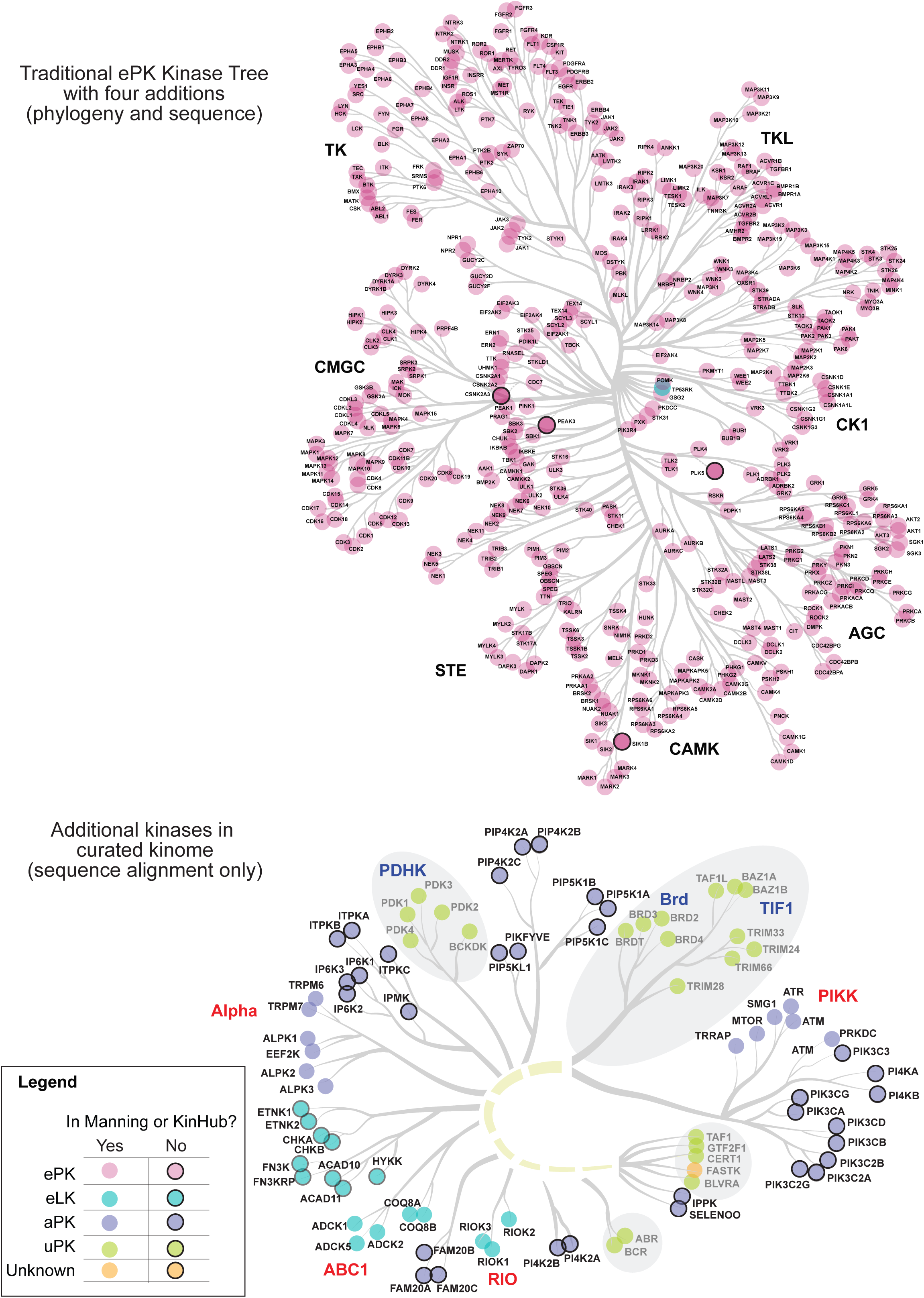
Two kinome trees for the curated kinome. Kinases from the curated kinome are visualized on the Coral kinase dendrogram (Metz et al., 2018) and a second kinome tree. Four ePKs previously omitted from the Coral kinase dendrogram (PLK5, PEAK3, CSNK2A3, and SIK1B) have been added to the main tree based on sequence alignment. The second small tree was created to depict relationships among eLKs with an aPK fold; it is based on sequence alignment of human proteins only. uPKs in the atypical kinase group (AKG) were excluded from the curated kinome, as described in the text, and are denoted by gray circles. Bolded black labels – kinase groups; blue bolded labels – uPK kinase families in AKG excluded from the curated kinome; red bolded labels – aPK or eLK kinase families in AKG included in the curated kinome. Note that the organization of the smaller tree does not imply the existence of established evolutionary relationship between eLK, aPK and uPKs (this is indicated by the use of dashes in the major branch of the dendogram).

The second tree aims to depict sequence similarity among kinases having eLK or aPK folds (such as the ethanolamine kinases and the choline kinases). AKG proteins having a uPK folds are included for reference, but are grayed out since they are not included in the curated kinome for reasons discussed above. It is important to note that the organization of this tree is based solely on sequence alignment and does not have extensive phylogenetic support. For this reason, the major branch (which links eLKs, aPKs and uPKs) is presented as a dashed line. Achieving greater confidence in evolutionary relationships may not be possible because sequence divergence in other organisms is too great. However, we judge the current depiction to be generally useful for studying the human kinome since many of the kinases depicted on the second tree are important disease genes and drug targets.

To explore the utility of the extended kinome for drug discovery and development, we analyzed a large-scale chemo-proteomic dataset collected using kinobeads (Klaeger et al., 2017). We found that 48 kinase domains that are part of the extended kinome, but not the curated kinome were bound to kinobeads and eight of these domains were competed-off by kinase inhibitors that are approved or in clinical trials. Hence, the extended kinome contains multiple enzymes such as pyridoxal kinase (PDXK) and adenosine kinase (ADK) that are missing from KinHub and Manning, but are relevant for establishing the specificity of kinase inhibitor drugs (**Figure S1B**) and may even be part of their mechanism of action (MoA). Moreover, non-protein kinases often participate in metabolic networks and protein kinases often function in signal transduction showing that the two are closely connected physiologically. We conclude that the extended and curated kinomes and their different subsets are useful in different settings.

### Consolidating kinase knowledge using an automatic literature reading engine

To compile existing knowledge about kinases and the biological networks they participate in we used the computational tool INDRA (the Integrated Network and Dynamical Reasoning Assembler, Box 1), (Gyori et al., 2017; Todorov et al., 2019). INDRA uses multiple natural language processing (NLP) systems (McDonald et al., 2016; Valenzuela-Escarcega et al., 2017) to extract computable statements about biological mechanisms from PubMed abstracts and full text articles in PubMedCentral. Unlike conventional bibliometric tools, INDRA aggregates data from a wide variety of databases (such as SIGNOR (the SIGnaling Network Open Resource) (Licata et al., 2020) and Pathway Commons (Cerami et al., 2011)) as well as information such as Target Affinity Spectra available via specialized resources such as the Small Molecule Suite (Target Affinity Spectra summarize known binding and not binding data for all compounds curated in ChEMBL (Moret et al., 2019). INDRA also normalizes and disambiguates biological entities from different sources and consolidates overlapping mechanistic interactions. The former process, known as “grounding,” involves the assignment of a unique identifier to all biological entities findable in text followed by linking the identifier to a reference database (for instance, HGNC and UniProt identifiers for human genes and proteins). Such normalization is particularly important because kinases are often described using multiple names (e.g., p38alpha, MAPK14, CSPB1, CSBP2, PRKM14, Mxi2, PRKM15 all refer to the same protein), and names often differ with the type of assay being performed. In such cases, the NLP systems that INDRA draws upon attempt to identify an optimal grounding, allowing INDRA to consolidate available mechanistic information for a given kinase with respect to phosphorylation sites, inhibitors, biological functions, etc. For example, the understudied kinase PEAK3 was originally known as C19orf35 and recently found to be a biologically active pseudokinase (Lopez et al., 2019). INDRA consolidates biological statements using PEAK3 and C19orf35 as synonyms and links them to the HGNC identifier 24793, and the now-standard HGNC name PEAK3. Another example involves CDK19, which was originally named CDK11 (confusingly, there are other CDK family members named CDK11A; HGNC:1730 and CDK11B; HGNC:1729). INDRA collected 50 mechanistic statements about CDK11 and another 50 statements about CDK19 and consolidated them under the standard HGNC name CDK19. Without INDRA’s internal homogenization scheme, half of the mechanistic information on CDK19 would have been lost to all but domain experts.

Whenever the information is available, INDRA statements are detailed with respect to molecular mechanism and they are linked to the underlying knowledge support (the database reference or citation, including database identifiers or specific sentences extracted from text and their PMIDs). For example, the INDRA network for the WEE2 understudied kinase (Figure 3A **and Figure S2A**) includes statements such as “*Phosphorylation (WEE2(), CDK1())*” and “*Inhibition (WEE2(), CDK1()).”* These machine and human-readable assertions imply that WEE2 is active in mediating an inhibitory phosphorylation event on CDK1 (Figure 3A). INDRA consolidates overlapping and redundant information: in many cases a single assertion has multiple pieces of evidence. For example, sentences from three PMID citations (PMIDs: 23287037, 21454751, and 19550110) were consolidated into one INDRA statement above: “*Phosphorylation (WEE2(), CDK1())*”. Each INDRA statement therefore represents a unique biological mechanism rather than a single mention in text. INDRA networks similar to the one shown for WEE2 were generated for all members of the curated kinome and can be visualized via the NDEx service (Pratt et al., 2015). A link to the NDEx network for each kinase is provided on kinome.org

**Figure 3.**
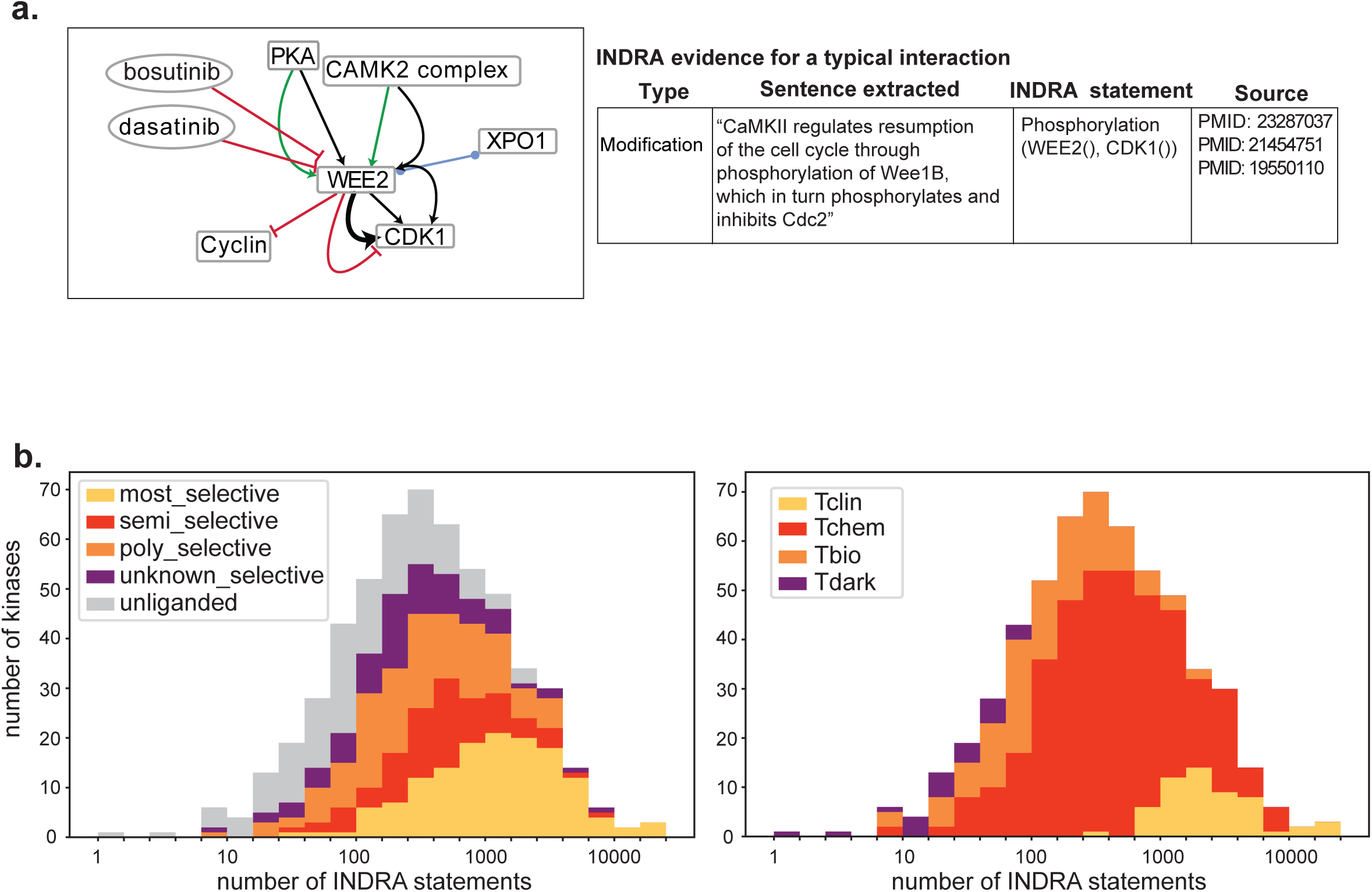
The composition of the understudied kinome. **(A)** Illustrative and simplified INDRA network automatically assembled for the WEE2 kinase. The table to the right shows the evidence extracted by INDRA for a single interaction (the bold arrow linking WEE2 and CDK1). An interactive version of this graph and a complete set of evidence can be found on NDEx (http://ndexbio.org). **(B)** Comparison of number of INDRA statements to levels of small molecule development. Left panel indicates selectivity level (MS-yellow, SS-red, PS-orange, UN-purple and grey-unliganded), right panel indicates drug development level according to PHAROS (Tclin-yellow, Tchem-red, Tbio-orange, Tdark-purple).

The number of INDRA statements extractable from text is a measure of the amount of knowledge available for each kinase. To compare the amount of knowledge available for each kinase in the curated kinome to the selectivity of chemical probes targeting that kinase we used selectivity information obtained from the Small Molecule Suite (Moret et al., 2019). This software tool mines diverse cheminformatic resources to determine which kinases are bound by small molecules in a most-selective (MS) and semi-selective (SS) fashion (Box 2). We also mined PHAROS (Nguyen et al., 2017), which classifies proteins based on whether or not they are bound by an FDA-approved drug (Tclin) or a tool compound (Tchem). We found that kinases that were more heavily studied were more likely to be classified as Tclin or Tchem in PHAROS and have inhibitors classified as MS or SS in Small Molecule Suite (Figure 3B). However, a substantial number of kinases with a high number of INDRA statements are bound only by relatively non-selective inhibitors, meaning that these kinases have a large body of knowledge, but development of selective drugs and tool compounds is trailing behind. Such kinases represent opportunities for development of new chemical probes. When we compared the number of INDRA statements to the TIN-X “novelty” score, which uses NLP of PubMed abstracts to assess the “novelty” and “disease importance” of a gene or protein (Cannon et al., 2017) we found that the two metrics were well correlated (Pearson’s correlation coefficient=0.81) **(Figure S2B).** Both metrics spanned ∼4 orders of magnitude from the least studied kinases (RSKR, with 0 INDRA statements) to the most (EGFR, with 24,856 INDRA statements). The distribution of INDRA statements on the curated kinome is also continuous and our use of the term “understudied” or “dark” kinome is based on an arbitrary cutoff corresponding to the least studied one-third of domains.

Using INDRA and TIN-X scores, we generated a new list of understudied kinases that is similar in scope to the IDG list, but based on objective criteria (schematized by the magenta box in **Figure S2B**). Starting with the curated kinome, this exercise yielded 182 least-studied kinase domains in 181 proteins of which 119 were part of the IDG set of kinases and 156 appear in Manning or KinHub. In the analyses that follows we use this computed set to define the “understudied kinome” (all sets are available at www.kinome.org under the “Knowledge” filter.) When the distribution of understudied kinases was viewed using the Coral tree depiction, an even distribution was observed across subfamilies, with the exception that only eight tyrosine kinases are relatively understudied (Figure 4). In many cases well-studied and understudied kinases are intermingled on the dendrogram (e.g., the CK1 subgroup) but in some cases an entire sub-branch is understudied (e.g., a branch with four TSSK and another with three STK32 kinases; dashed red outline). In yet other cases, a well-studied kinase is closely related to an understudied kinase, SIK1 and SIK1B or WEE1 and WEE2 for example, but it is unknown whether such pairs of isozymes are functionally redundant.

**Figure 4.**
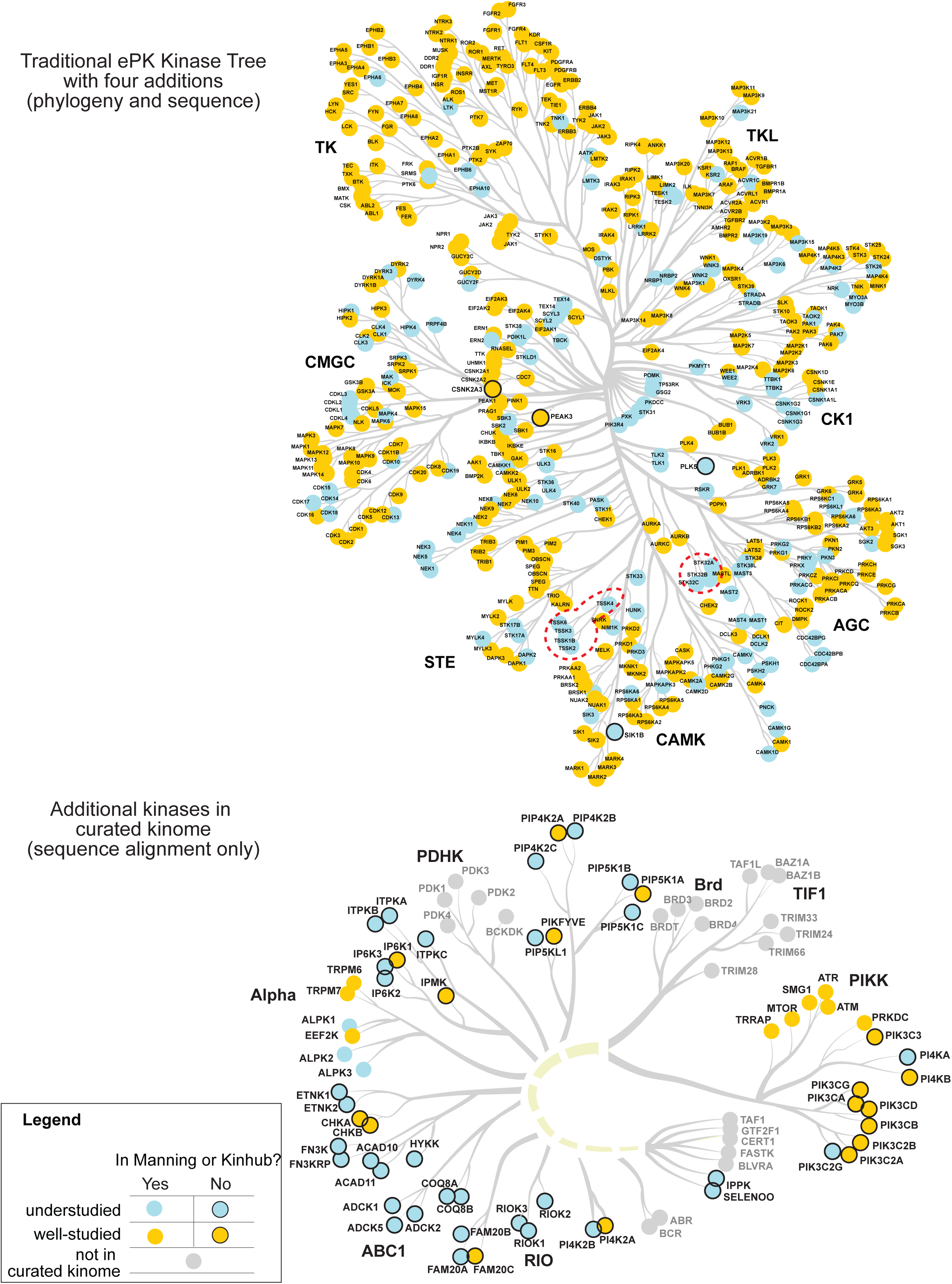
Understudied kinases on the Coral kinase dendrogram. Kinases from the curated kinome are visualized on the updated kinome trees (Figure 2) based on the Coral kinase dendrogram (Metz et al., 2018). The computed understudied kinome is shown in blue and well-studied kinases are shown in yellow. The atypical kinase group (AGC; denoted by a blue dashed line) as previously defined by Manning and KinHub as well as 46 aPK/eLK kinases in the curated kinome lies below the ePK tree. uPKs not in the curated kinome but in AKG are labelled in gray.

### Evidence for expression and function of understudied kinases

To consolidate existing data on the expression and possible functions of understudied kinases, we analyzed RNAseq data for 1019 cell-lines in the Cancer Cell Line Encyclopedia (CCLE) (Nusinow et al., 2020), proteomic data for 375 cell-lines in the CCLE (Nusinow et al., 2020) and loss of function data in the Cancer Dependency Map (DepMap). The DepMap was generated using lentivirus-based RNAi or CRISPR/Cas9 libraries in pooled screens to identify genes that are essential in one or more of ∼1000 cell-lines (Tsherniak et al., 2017).

Based on RNASeq data, both well-studied and understudied kinases were observed to vary substantially in abundance across 1019 CCLE cell-lines. Using the common threshold of RPKM ≥1 (Reads Per Kilobase of transcript, per Million mapped reads) (Kryuchkova-Mostacci and Robinson-Rechavi, 2017) evidence of expression was found in at least one cell-line for 176 out of 181 understudied kinases (Figure 5A). Some understudied kinases were as highly expressed as well-studied kinases: for example, NRBP1 and PAN3 and the lipid kinases PI4KA and PIP4K2C all had maximum expression levels similar to that of the abundant and well-studied LCK tyrosine kinase. Overall, however, understudied kinases had significantly lower maximum mRNA expression levels than well-studied kinases by multiple criteria (2.1 vs 5.8 RPKM median expression level, p-value=4.6×10^-8^; 36 vs 71 RPKM maximum expression level, p-value=2.2×10^-16^ by Wilcoxon rank sum test). In CCLE proteomic data we observed that 367 kinases from the curated kinome were detected at the level of at least one peptide per protein; 110 of these were understudied kinases. The difference between proteomic and mRNA data is likely to reflect the lower sensitivity of shotgun proteomics, but some kinases might also be subjected to translational regulation.

**Figure 5.**
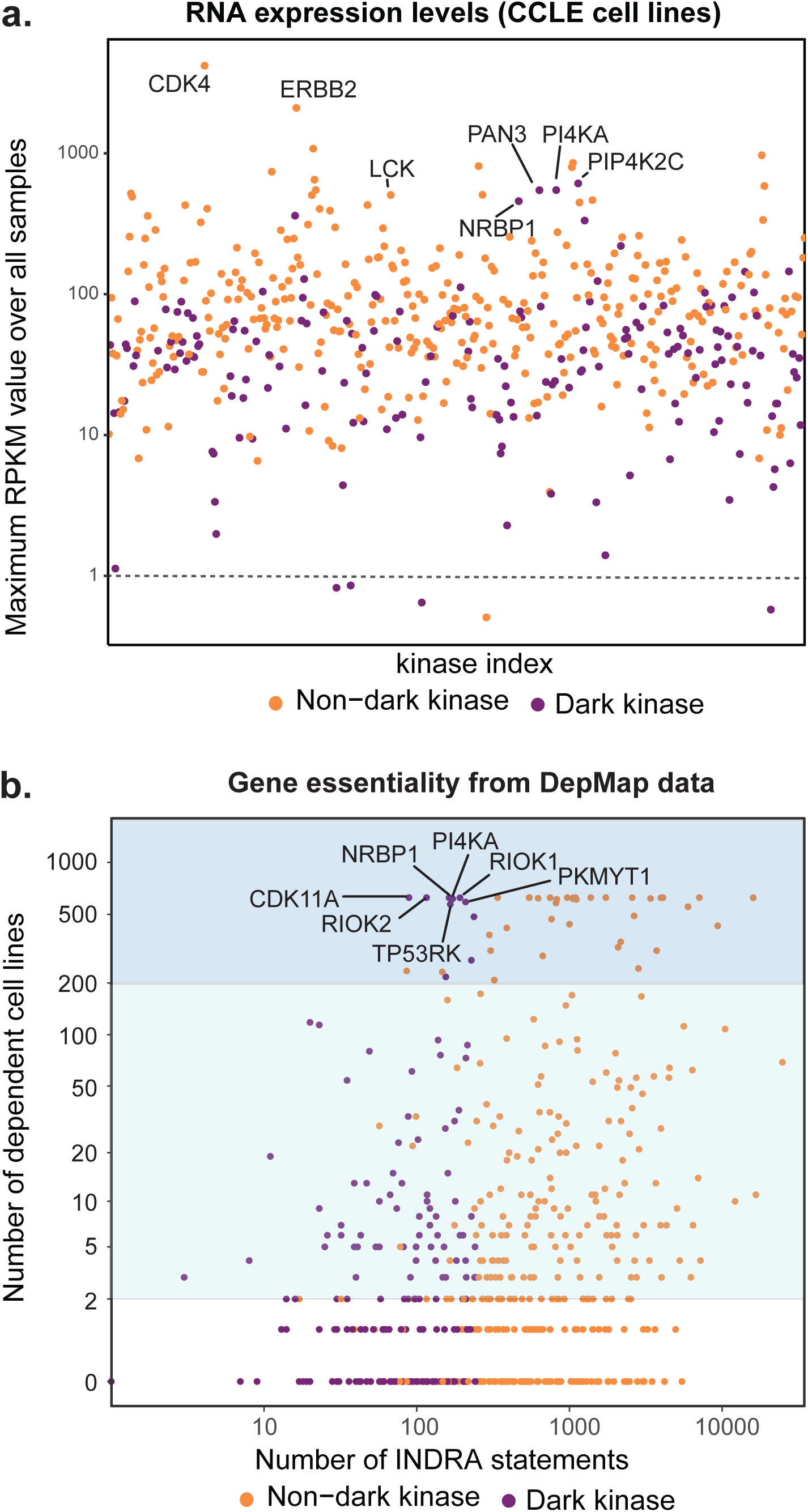
Evidence for understudied kinase expression and function. **(A)** Maximum expression level (RPKM value) for each gene in the understudied kinome list across 1039 cell-lines curated in the CCLE database (Barretina et al., 2012). Understudied kinases are colored in purple, well-studied kinases in orange. Dotted line indicates a RPKM threshold of 1, above which genes were designated as “expressed” based on an established metric (Kryuchkova-Mostacci and Robinson-Rechavi, 2017). **(B)** Number of cell-lines for which the Dependency Map (Tsherniak et al., 2017) score indicates essentiality (using the recommended *Post-Ceres* (Meyers et al., 2017) value of ≤ −0.5). Understudied kinases are colored in purple, well-studied kinases in orange; HGNC symbols for genes essential in all cells in the Dependency Map are shown. Blue shading denotes genes essential in one-third or more of cell-lines and yellow denotes genes essential in two or more lines.

Analysis of DepMap data showed that 10 understudied kinases are essential in at least one third of the 625 cell-lines tested to date (Figure 5B; dark blue shading), and 88 understudied kinases are essential in at least two lines (light blue shading). We conclude that a substantial number of understudied kinases are expressed in human cell-lines and a subset are required for cell growth. These data are likely to underestimate the breadth of kinase expression and function: proteins can impact cellular physiology when expressed at low levels (not detectable by shotgun proteomics) and genes can have important functions without necessarily resulting in growth defects assayable by the DepMap methodology.

### Understudied kinases in disease

To study possible roles for understudied kinases in pathophysiology, we mined associations between diseases and either gene mutations or changes in gene expression. We mined The Cancer Genome Atlas (TCGA), Accelerating Medicines Partnership - Alzheimer’s Disease (AMP-AD) dataset and a microarray dataset on changes in gene expression associated with chronic obstructive pulmonary disease (COPD; (Rogers et al., 2019)). COPD progressively impairs a patient’s ability to breathe and is the third leading cause of death in the US (National Center for Health Statistics (US), 2016). In TCGA, we compared the frequency of mutations between understudied and well-studied kinases under the assumption that the two sets of kinases are characterized by the same ratio of passenger to driver mutations (Garraway and Lander, 2013) and we then looked for differential RNA expression relative to matched normal tissue (Figure 6A). In line with most TCGA analyses, mutations and differential expression were scored at the level of genes and not domains; thus, observed mutations may affect functions other than kinase activity. We performed differential expression and mutation frequency analyses for individual tumor types and for all cancers as a set (the PanCan set). With respect to differential gene expression, we found that understudied and well-studied kinases are equally likely to be over or under-expressed in both PanCan data and in data for specific types of cancer (in a Rank-sum two-sided test with H_0_ = well-studied and understudied kinases have similar aberrations, p=0.86) (Figure 6). For example, in colorectal adenocarcinoma, the understudied kinase MAPK4 is one of the three most highly downregulated kinases whereas LMTK3, NEK5 and STK31 represent four of the seven most highly upregulated kinases (**Figure S3A**). This is consistent with a report that overexpression of STK31 can inhibit the differentiation of colorectal cancer cells (Fok et al., 2012).

**Figure 6.**
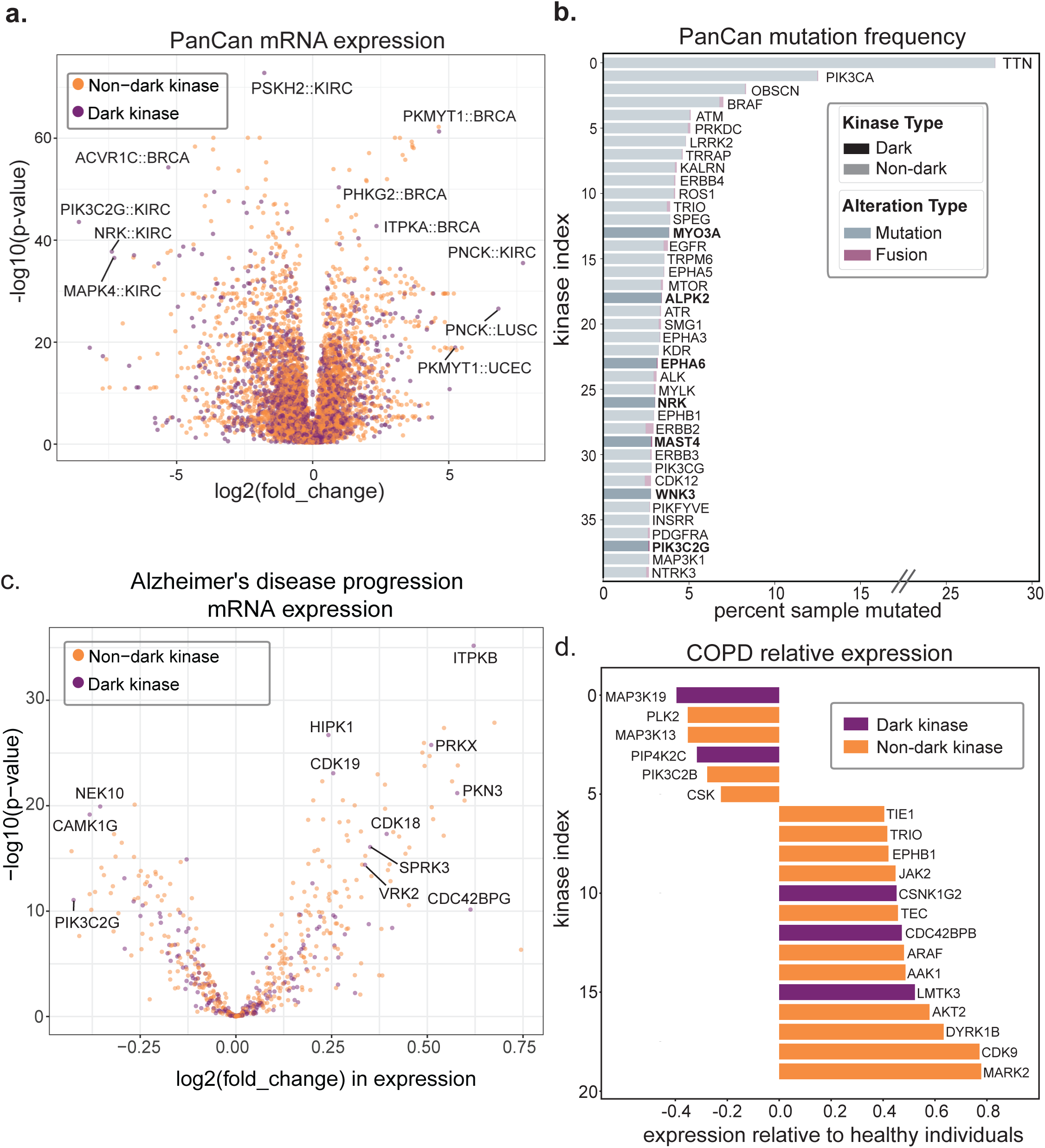
Understudied kinases in diseases. Data on differential expression, mutation, amplification or deletion of genes containing domains from the curated human kinome in disease databases. Understudied kinases are colored in purple, well-studied kinases in orange. **(A)** Pan-cancer (PanCan) differential mRNA expression for both the understudied and the well-studied kinases based on data in TCGA and accessed via cBioPortal (Cerami et al., 2012). No significant difference between the understudied and well-studied kinome was observed with respect to the frequency of differential expression relative to matched normal tissue. HGNC symbols and cancer type abbreviations for selected outlier genes and diseases are shown. BRCA - Breast invasive carcinoma; KIRC - Kidney renal clear cell carcinoma; LUSC - Lung squamous cell carcinoma; UCEC - Uterine Corpus Endometrial Carcinoma. **(B)** Mutation frequency for most frequently mutated kinases in PanCan. Understudied kinases are shown in solid color; well-studied kinase in transparent color. Fusion-mutations are shown in magenta. HGNC symbols are displayed next to each bar with bold denoting understudied kinases. **(C)** Differential gene expression in early versus late stage Alzheimer’s disease. Samples were aged matched prior to calculation of differential expression values. HGNC symbols are shown for outliers displayed. **(D)** Relative gene expression levels in COPD versus healthy individuals. Kinases are sorted by their relative expression. HGNC symbols are displayed next to each bar with purple denoting understudied kinases and orange denoting well-studied kinases.

By mining PanCan, we also found that five understudied kinases were among the 30 most frequently mutated human kinases; for example, the ∼3% mutation frequency of the understudied MYO3A kinase is similar to that of the oncogenic RTKs EGFR and ERBB4 (but lower than the ∼12% mutation frequency for the lipid kinase PIK3CA) (Figure 6B). In diffuse large B-cell lymphoma (DLBCL), the understudied kinase ITPKB, which has been reported to phosphorylate inositol 1,4,5-trisphosphate and regulate B cell survival (Schurmans et al., 2011), is more frequently mutated than KDR (∼13% vs. 8% of patient samples, **Figure S3B**). Overexpression of KDR is known to promote angiogenesis and correlate with poor patient survival (Gratzinger et al., 2010; Holmes et al., 2007; Jørgensen et al., 2009). More recently a case study reported that patients carrying an ITPKB C873F substitution mutation had Richter’s syndrome (which is characterized by the sudden development of B-cell chronic lymphocytic leukemia into the faster-growing and more aggressive DLBCL) and clones harboring the ITPKB C873F mutation exhibited higher growth rates (Landau et al., 2017). Recurrent mutation, over-expression and under-expression in TCGA data is not evidence of biological significance *per se*, but systematic analysis of TCGA data has been remarkably successful in identifying genes involved in cancer initiation, progression, and drug resistance. Our analysis shows that understudied kinases are nearly as likely to be mutated or differentially expressed in human cancer as their well-studied kinase homologues, making them good candidates for future testing as oncogenes or tumor suppressors.

The AMP-AD program Target Discovery and Preclinical Validation (Hodes and Buckholtz, 2016) program aims to identify molecular features of AD at different disease stages. We compared mRNA expression at the earliest stages of AD to late-stage disease in age matched samples (Figure 6C) and found that the understudied kinases ITPKB and PKN3 were among the five most upregulated kinases while NEK10 was substantially downregulated. A similar analysis was performed for COPD, based on a study by Rogers et al (Rogers et al., 2019) of five COPD microarray datasets from Gene Expression Omnibus (GEO) and two COPD datasets from ArrayExpress that aimed to identify genes with significant differential expression in COPD. By comparing the expression of genes in COPD patients to gene expression in healthy individuals, Rogers et al. identified genes significantly up and down regulated in patients (adjusted p-value < 0.05). We found that the understudied kinase PIP4K2C, which is potentially immune regulating (Shim et al., 2016), was significantly downregulated in individuals with COPD (adjusted p-value = 0.048). Additionally, CDC42BPB, which is possibly involved in cytoskeleton organization and cell migration (Tan et al., 2008, 2011), was significantly upregulated (adjusted p-value = 0.026) (Figure 6D). In total, five understudied kinases versus fifteen well-studied kinases were differentially expressed in COPD patients. All kinases with biological relevance as implicated by analysis of DepMap, TCGA, AMPAD, and COPD datasets can be found at kinome.org under the “Biological Relevance” tab. As additional data on gene expression and mutation become available for other diseases, it will be possible to further expand the list of understudied kinases potentially implicated in human pathophysiology.

### An understudied kinase network regulating the cell cycle

Inspection of INDRA networks showed that understudied kinases, like well-studied ones, function in networks of interacting kinases. One illustrative example involves control of the central regulator of cell cycle progression, CDK1, by the understudied kinases PKMYT1, WEE2, BRSK1 and NIM1K (Figure 7). Although the homologues of some of these kinases have been well studied in fission and budding yeast, less is known about the human kinases (Wu and Russell, 1993). WEE2, whose expression is described to be oocyte-specific (Sang et al., 2018) but can also be detected in seven CCLE lines, six from lung cancer and one from large intestine, is likely to be similar in function to the well-studied and widely-expressed homologue WEE1, which phosphorylates CDK1 on the negative regulatory site T15 (Sang et al., 2018). The related kinase PKMYT1 phosphorylates CDK1 on the Y14 site to complete the inhibition of CDK1 (Liu et al., 1997; Mueller et al., 1995). These modifications are removed by the CDC25 phosphatase, which promotes cell cycle progress from G2 into M phase (Santamaría et al., 2007). PKMYT1 and WEE1 are essential in nearly all cells, according to DepMap (DepMap, 2019) (although WEE2 is not). Upstream of WEE1, the relatively understudied kinases BRSK1 (127 INDRA statements) and NIM1K (27 INDRA statements) and the related kinase BRSK2 (176 statements) function to regulate WEE1. Neither PKMYT1, BRSK1 nor NIM1K have selective small molecule inhibitors described in the public literature (Asquith et al., 2020); several WEE1 inhibitors are in clinical development however (Matheson et al., 2016), and these molecules are likely to inhibit WEE2 as well. It is remarkable that enzymes so closely associated with the essential cell cycle regulator CDK1, remain relatively understudied in humans (Wu and Russell, 1993). This is particularly true of PKMYT1 and NIM1K which are frequently upregulated in TCGA data.

**Figure 7.**
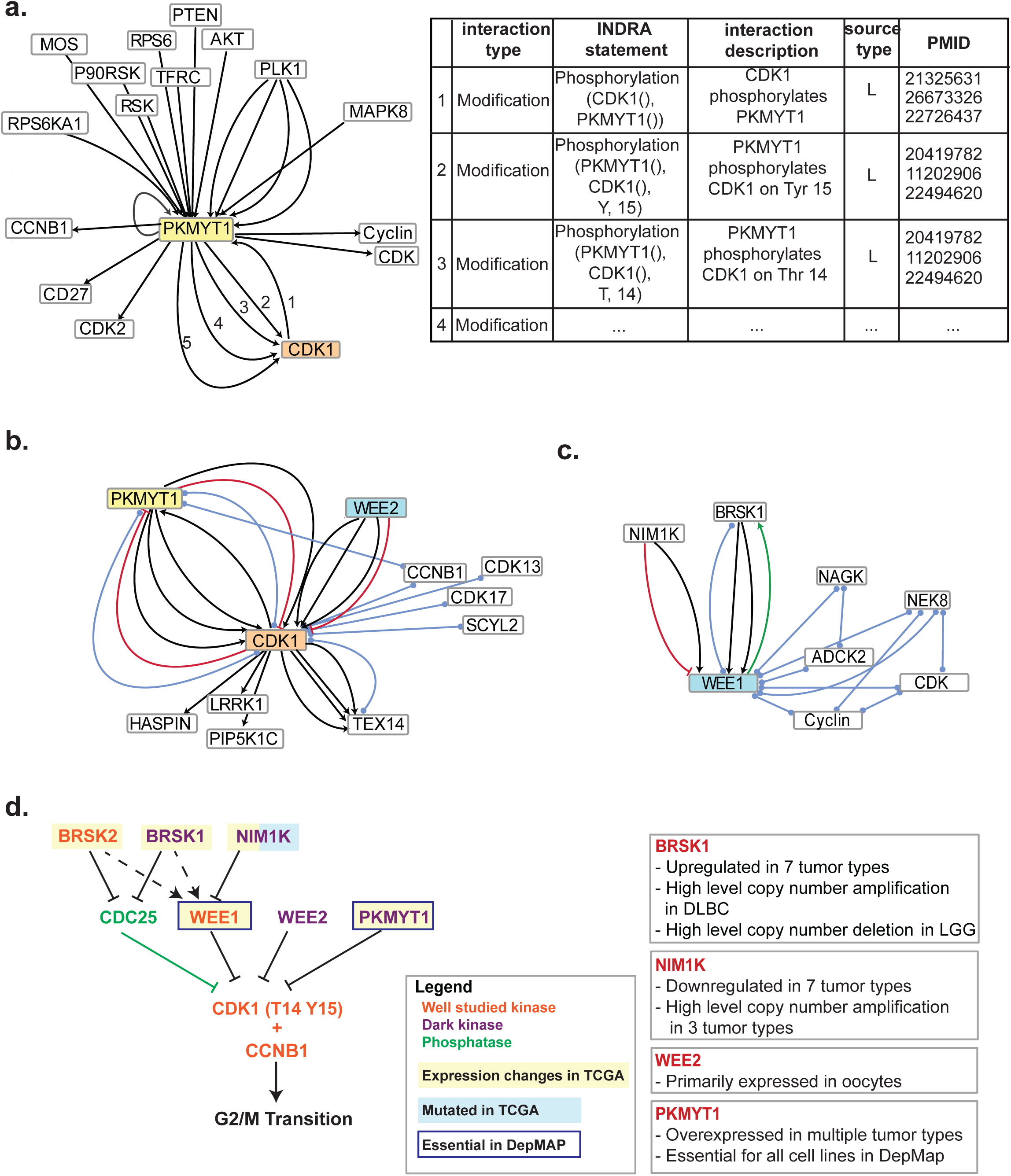
An understudied kinase network regulating the cell cycle. **(A)** A partial network (left) and statement table (right) for proteins interacting with PKMYT1 according to the INDRA database. The source “literature” is denoted by “L”. **(B)** INDRA network for CDK1 showing interacting understudied kinases. Black arrows denote protein modifications; blue lines denote complex formation; red arrows denote inhibition; green arrows denote activation. **(C)** INDRA network for WEE1 showing interacting understudied kinases. Color code is the same as in panel B. **(D)** Manually curated signaling network based on known regulatory mechanisms for PKMYT1, CDK1 and WEE1. The network comprises four understudied kinases (BRSK1, NIM1K, WEE2 and PKMYT1), three well-studied kinases (BRSK2, WEE1, CDK1), and the protein phosphatase CDC25.

### Inhibition of understudied kinases by approved drugs

Kinase inhibitors, including those in clinical development or approved as therapeutic drugs, often bind multiple targets (Klaeger et al., 2017). We therefore asked whether understudied kinases are targets of investigational and FDA-approved small molecules by using the Small Molecule Suite (Moret et al., 2019) to mine public data for evidence of known binding and known not-binding. We identified 17 understudied kinases that might be inhibitable with MS or SS selectivity by approved drugs or compounds that have entered human clinical trials (some of which may not currently be in active development). For example, the approved anti-cancer drug sunitinib is described in the Small Molecule Suite as binding to the understudied kinases STK17A, PHGK1 and PHGK2 with binding constants of 1 nM, 5.5 nM and 5.9 nM respectively based on curated dose-response data from enzymatic assays (Davis et al., 2011; Fabian et al., 2005; Karaman et al., 2008; McDermott et al., 2007; Zarrinkar et al., 2009) (**Figure S4, Table S3**). This contrasts with affinities of 30 nM to 1 µM for VEGF receptors (the KDR, FLT1 and FLT4 kinases) and 200 nM for PDGFRA, which are well established and likely clinically relevant targets for sunitinib. To validate binding to understudied kinases predicted from Target Affinity Spectra (Moret et al., 2019) we performed dose-response experiments using a commercial panel of enzymatic phospho-transfer assays (the Reaction Biology kinase assay panel, www.reactionbiology.com, Malvern, PA) that includes 91 understudied kinases. We confirmed 19 cases in which a clinical-grade compound inhibited an understudied kinase with a binding constant of 1 µM or less (Figure 8A). In six cases, the drug was as or more potent on an understudied kinase than on a target described as part of the drug’s mechanism of action (the nominal target; Box 2). In two cases, predicted binding could not be confirmed experimentally (inhibition of LTK by crizotinib and inhibition of CDK19 by linifanib; denoted by asterisk in Figure S6). Moreover, two understudied kinases do not have available assays (STK17B and CDK11A; denoted with a double asterisk in Figure S6) and it was not possible to assay their activities. Follow-on biochemical and functional experiments will be required to determine if understudied kinases actually play a role in the therapeutic mechanisms of these and other approved drugs.

**Figure 8.**
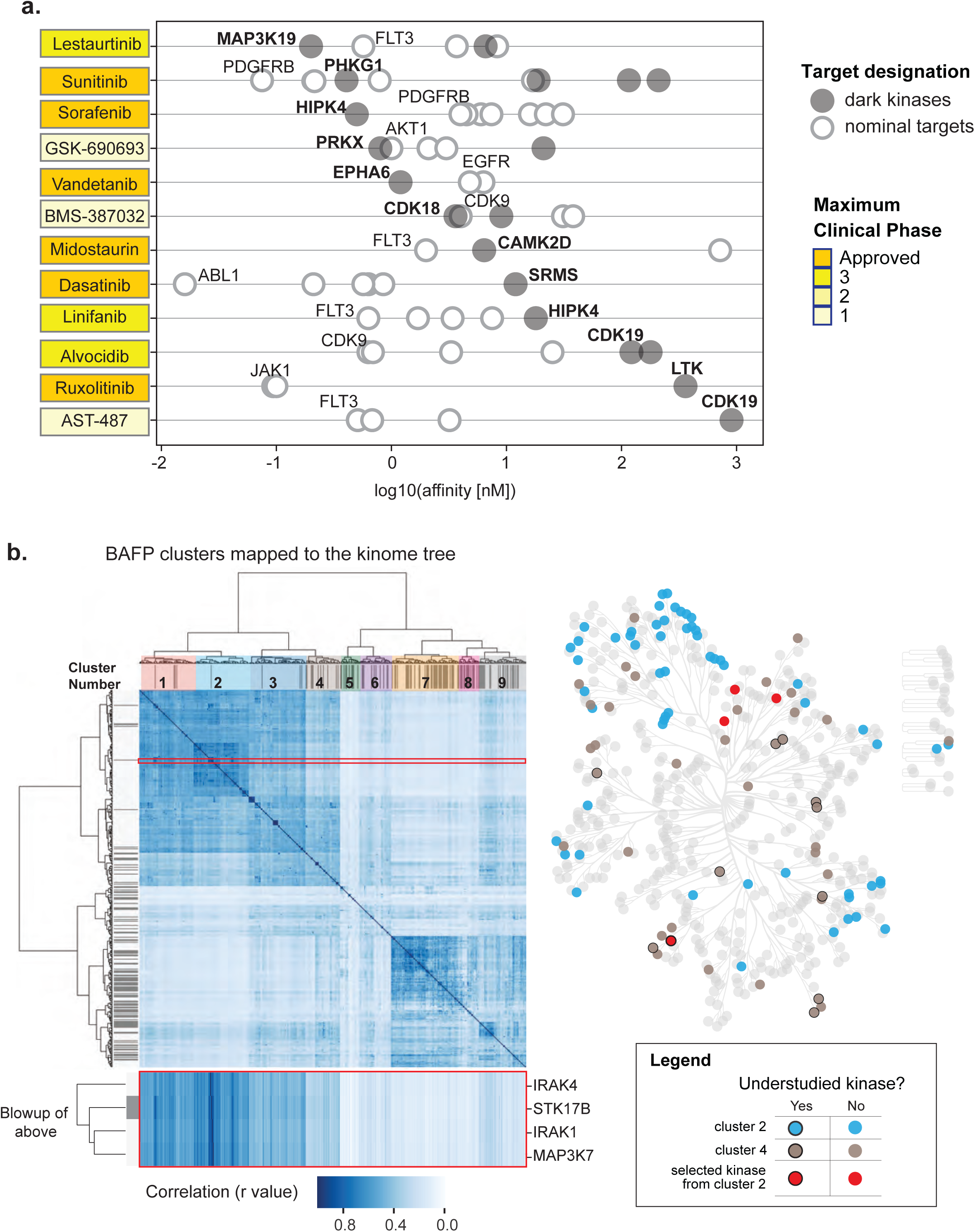
Inhibition of understudied kinases by clinical grade compounds and approved drugs. **(A)** Binding constants of clinical compounds on their nominal targets and dark kinases. Dark kinases described in the Small Molecule Suite to interact with the clinical compounds were confirmed using enzymatic assays. The nominal targets of these clinical compounds and their corresponding binding affinities were extracted from the National Center for Advancing Translational Sciences (NCATS) Inxight: Drugs portal (https://drugs.ncats.io/). Dark circle: binding constants of clinical compounds on dark kinases; white circle: binding constants of clinical compounds on their nominal targets. **B)** Clustering of understudied kinases by Bayes Affinity Fingerprint (BAFP) – a measure of the shape of binding pocket. Dark lines in the margin denote understudied kinases. A blowup of BAFP values for four kinases (red box), one of which is understudied (STK17B), is shown below. To the right, cluster 2 (blue) and cluster 4 (brown) from BAFP clustering are projected onto the Coral dendrogram. Four selected kinases from cluster 2 (in the red box on the left) are colored red.

The potential for development of new compounds that inhibit understudied kinases based on modification of existing kinase inhibitors can be assessed in part by examining the structures of kinase binding pockets using Bayes Affinity Fingerprints (BAFP) (Box 3) (Bender et al., 2006; Nguyen et al., 2013). In this cheminformatics approach, each small molecule in a library is computationally decomposed into a series of fragments using a procedure known as fingerprinting. The conditional probability that a compound binds to a specific target (as measured experimentally in profiling or enzymatic assays) given the presence of a chemical fragment is then calculated. Each target (protein) is thereby associated with a vector comprising conditional probabilities for binding to fragments found in the fingerprints of compounds in the library. Subsequently, the correlation of conditional probability vectors for two proteins is used to evaluate similarity in binding pockets from the perspective of a chemical probe. BAFP vectors were obtained from a dataset of ∼5 x 10^6^ small molecules and 3 x 10^3^ targets for which known binding and non-binding data are available from activity profiling.

We found that the majority of kinase domains fell in two clusters based on their BAFP vectors; both clusters had multiple understudied and well-studied kinases. The high similarity of the two sets of kinases in “compound binding space” suggests that many more kinase inhibitors than those described in Figure 8A may bind understudied kinases or could be modified to do so (Figure 8B**, Figure S5**). For example, the clustering of IRAK1, IRAK4, STK17B and MAP3K7 by BAFP correlation (highlighted in Figure 8B) shows that the STK17B binding pocket is likely very similar to that of IRAK1, IRAK4 and MAP3K7 and that compounds binding these well-studied kinases, such as lestaurtinib and tamatinib may also bind STK17B. Based on this, it may be possible to design new chemical probes with enhanced selectivity for STK17B by starting with compound series based on lestaurtinib or tamatinib.

Other useful tools for development of new small molecule probes are commercially available activity assays and experimentally determined NMR or crystallographic protein structures. Of 181 understudied kinases, 101 can currently be assayed using the popular KINOMEscan platform (Fabian et al., 2005); and 74 are found in both Reaction Biology and KINOMEscan. Since the Reaction Biology assay measures phospho-transfer activity onto a peptide substrate, the availability of an assay provides further evidence that at least the 91 understudied kinases in this panel are catalytically active. Searching the Protein Data Bank (PDB) reveals that 53 understudied kinases have at least one experimentally determined structure (for the kinase domain at least). HASPIN has 18 structures, the highest of all understudied kinases, followed by PIP4K2B, CLK3, and CSNK1G3 (14, 10 and 10 structures, respectively) (**Figure S6, Table S2**). Many of these structures were determined as part of the Protein Structure Initiative (Burley et al., 2008) and its successors but have not been subsequently discussed in the published literature. The availability of these resources and data are summarized in Supplementary Table 1 and available on www.kinome.org under the “Resources” tab.

### A small molecule library to perturb the kinome

To create an optimal set of chemical probes for the well-studied and understudied kinomes we assembled the ‘OptimalKinase Library v2.0’ (OKL2.0) inhibitor library. A similar library was proposed in previous work (OKL-v1.0 (Moret et al., 2019)), but we subsequently found that several of the inhibitors in that library were not commercially available and were expensive to synthesize. In OKL2.0 we include only small molecules available from commercial sources and also account for the updated kinome membership described in this paper.

The OKL2.0 library contains 194 compounds in total which are selected so that each kinase is ideally bound by two small molecules having the highest possible selectivity and as great a difference in chemical structure as possible (Moret et al., 2019). The library also includes approved drugs or compounds in clinical trials that inhibit a kinase with an affinity less than 1 µM; in many but all cases, these are also highly selective compounds. Not all kinase domains can be bound with MS selectivity: using compounds in OKL2.0, 127 domains in the curated kinome are bound with MS and 74 with SS selectivity (as compared to 136 MS and 76 SS compounds in OKL-v1.0)(Figures 9A,B **and S7A**). As additional measures of the quality of the OKL2.0 library we confirmed low correlation in chemical structure and diversity in Target Affinity Spectra (TAS) and phenotypic fingerprints (Figure 9C**; S7B-D**).

**Figure 9.**
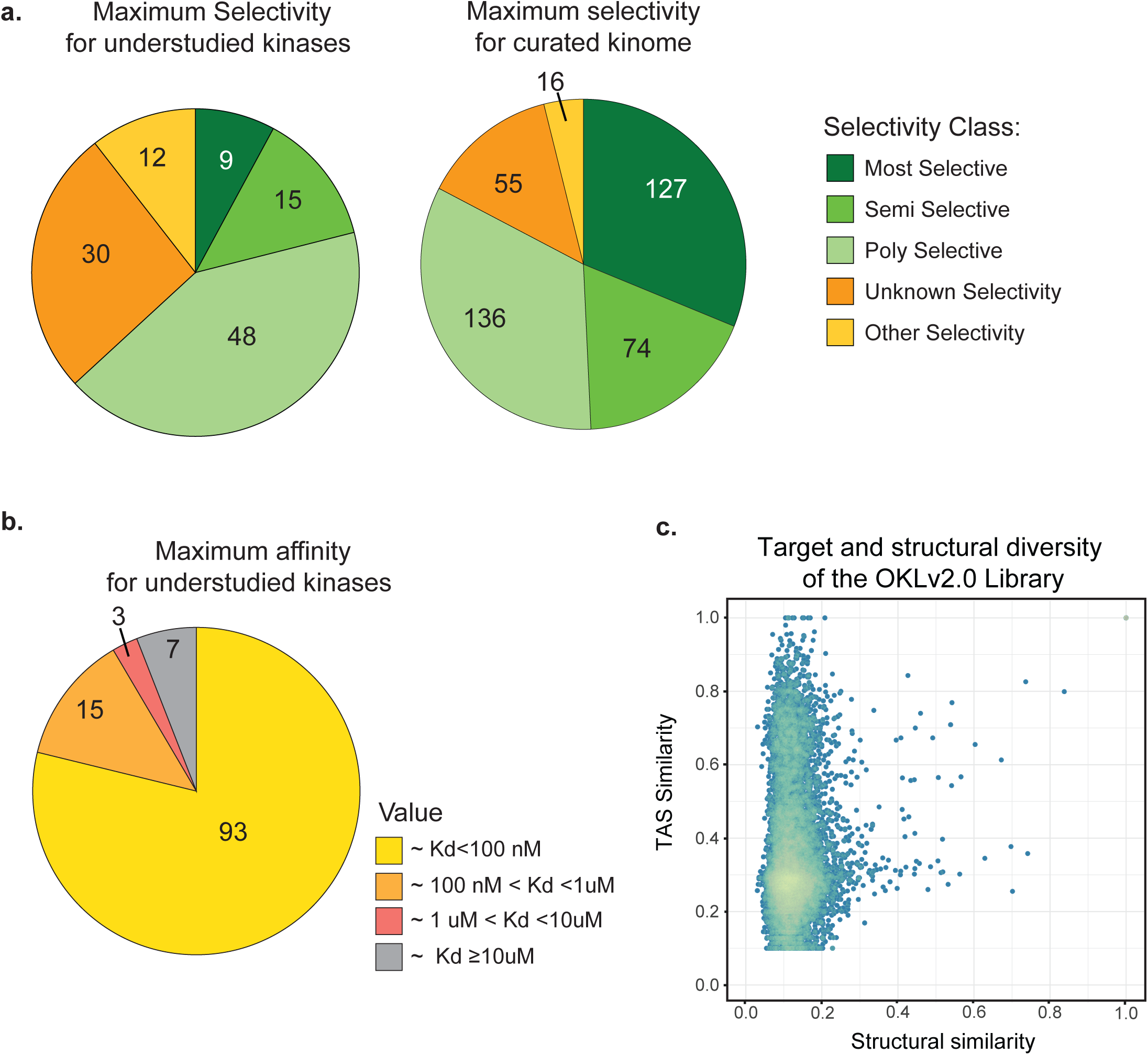
Performance of the OKL2.0 library for the understudied kinome. **(A)** Maximum selectivity per kinase domain achievable with compounds in OKL2.0. 114 of 182 members of the understudied kinome are shown to the left and 408 of 557 members of the curated kinome on the right. Legend on the right of the pie chart explains the color scheme. **(B)** Maximum affinity (lowest Kd) achieved by compounds in OKL2.0. Legend on the right of the pie chart explains the color scheme. **(C)** Plot of the similarity in Target Affinity Spectra versus structural similarity based on the Tanimoto score of Morgan2 fingerprints.

Focusing on the 182 understudied kinases, 9 can be inhibited with MS selectivity and 15 with SS selectivity. When we examined Target Affinity Spectra and considered only binding affinity not selectivity, we found that 93 of 182 understudied kinases could be bound by at least one compound in the OKLv2.0 with Kd <100 nM (18 more had lower affinity chemical ligands: 15 bound at 100 nM <Kd <1 µM, and 3 at 1 µM, <Kd <10 µM). These ligands are not likely to be useful as chemical probes (Arrowsmith et al., 2015) but they may serve as starting points for the development of new, more selective compounds. We conclude that the OKL2.0 library has optimal qualities given the current state of public knowledge for perturbing both the well-studied and understudied kinomes.

## TEXT BOX ONE

### Using INDRA to consolidate mechanistic knowledge

INDRA (Integrated Network and Dynamical Reasoning Assembler) is an open-source software system for automatically collecting information about biological entities (primarily proteins, small molecules, and biological processes) and then assembling this information into mechanistic models and networks. INDRA uses several different natural language processing (NLP) systems, each with their own strengths and weaknesses, to extract relationships between biological entities from text in a computable form, primarily from published papers. INDRA also integrates structured and semi-structured information from pathway databases and chemical biology resources.

INDRA converts diverse sources of information into standardized machine interpretable mechanistic assertions known as “INDRA Statements.” Each biological entity is assigned identifiers in one or more reference databases (HGNC, UniProt, ChEBI, GO, etc.) so that multiple names referring to the same entity are standardized and knowledge of the biological entity is preserved even if its name changes. This is particularly important in the case of kinases for which both standardized and informal naming schemes co-exist. “INDRA Statements” further normalize the variability in describing biochemical reactions by mapping the extracted mechanism to a standardized template. Redundant information from multiple sources is consolidated this way into a single assertion. For example, a phosphorylation event might be described at different levels of specificity in different texts (with or without the identity of a modified residue, or the presence of a bound entity), and INDRA will combine sources of information into the most precise description for which evidence is available. INDRA Statements keep track of the provenance of the underlying evidence for a statement in the form of PubMed IDs or database entry identifiers. Thus, each INDRA Statement represents all the information about a particular biochemical mechanism that can reliably be combined.

## TEXT BOX 2

### Targets and selectivity of small molecule drugs

Small molecule drugs are rarely specific for a single target protein. We and others (Moret et al., 2019; Wang et al., 2016) have therefore developed methodology to quantify and account for this polypharmacology (the binding of a small molecule to multiple target proteins). Typically, small molecules are designed with a single protein in mind and this usually becomes the target that is most associated with that compound (we refer to this as the “nominal target”). For approved drugs, the nominal target is often found on the FDA label but it is not necessarily the highest affinity target or the only target involved in therapeutic mechanism of action. It is possible to account for data on known binding and non-binding of small molecules to multiple target using the *Target Affinity Spectra* (TAS) and the *selectivity score*. TAS is a vector that summarizes binding information from a variety of sources (e.g, dose-response data, one-dose percent inhibition data and binding assessments in scientific literature) by assigning an assertion score of 1, 2 or 3 for very-tight to less-tight binding and 10 for confirmed non-binding. The selectivity score is a measure assigned to compound-target binding interactions for which dose-response data are available. A compound that has three known target-interactions (from a TAS vector) will get three selectivity scores, one for each interaction. The selectivity score is calculated from available data on the absolute binding affinity, differential “on target” affinity as compared to the “off-target affinity”, the *p-value* between the distributions for “on” and “off” targets, and “research bias.” Research bias accounts for differences in the amount of binding data that is available; in the absence of a bias estimate, a poorly studied compound can appear more selective than a well-studied one simply because fewer targets have been tested and fewer “off-targets” identified. The MS assertion is assigned to compounds that have an absolute affinity <100 nM and for which the on-target Kd (more precisely, the first quartile of available dose-response data for that target) are > 100 lower than the Kd of all other targets (based on the first quartile of data available for them), p-value of ≤ 0.1, and research bias ≤ 0.2 (see (Moret et al., 2019) for details). The SS assertion is about 10-fold less strict with regard to absolute and differential affinity (see methods).

## TEXT BOX 3

### Bayesian Affinity Fingerprints for the assessment of binding-pocket similarity

Fingerprinting is an approach widely used to calculating the structural similarity of sets of small molecules by computationally decomposing them into small fragments. When a compound library is fingerprinted, a set of unique fragments is created, making it possible to represent each parent compound as a vector of zeros and ones specifying which fragments it contains. The fingerprints can then be compared using standard vector calculus or machine learning approaches. Bayesian Affinity Fingerprints (BAFP) (Bender et al., 2006) are a special type of fingerprint in which each fragment is annotated with a conditional probability that it binds to a target of interest (typically with a 10 µM cutoff). A large dataset on compound binding comprising many compounds and targets and their respective affinities (as measured using biochemical dose-response assays) is required to calculate these conditional probabilities. The product of Bayesian fingerprinting is a matrix with fragments on one axis (columns) and protein-targets on the other axis (rows). Each number in the matrix is the conditional probability that a particular compound binds to the target when the fragment is part of that compound (i.e., the fragment is in its fingerprint). This matrix can be used to describe protein-targets as a vector of conditional probabilities that a small molecule will bind given the presence of a particular chemical fragment. The correlation between two vectors can also be used as a measure of the similarity between binding pockets. For example, if two targets A and B bind similar chemical structures (i.e., have highly correlated BAFPs) then a compound that bind protein A can be expected to bind protein B as well.

## DISCUSSION

In this paper we mine sequence and structural information to provide complementary definitions of proteins comprising the human kinome. We also use NLP and knowledge assembly to quantify available information about individual kinase domains and summarize the data in a web resource (www.kinome.org) intended for use in systematic functional genomics, chemical biology and drug discovery. This resource is updated regularly and can be used to generate more or less expansive sets of kinases in a data-driven manner based on criteria such as function, sequence homology, known or predicted structure. We find that the amount of published information available on each kinase varies ∼10^4^-fold from the best-studied drug targets, such as SRC and EGFR, to less studied kinases such as PKMYT1 that are essential for cell growth and overexpressed in diseases such as cancer. The related web resource <u>darkkinome.org</u> (Berginski et al., 2020) aggregates information and INDRA networks on the understudied one-third of the kinome and is intended to motivate further study of these proteins. We find that existing chemical matter - including some clinical grade compounds – are available to inhibit several understudied kinases with at least partial selectivity, providing new opportunities for development of novel chemical probes and perhaps also therapeutic drugs.

Determining the set of proteins that comprise the human kinome is not a trivial task because multiple overlapping but non-identical definitions exist, often based on different and ambiguous molecular or functional criteria. This is not simply a matter of semantics, because the full substrate specificity of many kinases remains unknown from experimental studies, particularly in the case of understudied kinases. For example, it is increasingly clear that some kinases can phosphorylate both small molecules and proteins (Yang et al., 2012). Cleanly separating protein kinases from lipid and some metabolite kinases may not be possible, and it may not make sense from a biological perspective: PI3K, for example, is commonly described as being part of the AKT/mTOR kinase cascade, even though the PI3K proteins are inositol lipid kinases and AKT/mTOR are protein kinases.

The human kinome includes ∼50 so-called “pseudokinases” that, based on sequence alignment, lack one or more residues involved in catalytic activity. These residues include the ATP-binding lysine (K) within the VAIK motif, the catalytic aspartate (D) within the HRD motif and the magnesium binding D within the DFG motif (Kwon et al., 2019). Nonetheless, some pseudokinases have important functions in signal transduction. For example, the EGFR family member ERBB3/HER3 is a pseudokinase that, when bound to ERBB2/HER2, forms an active receptor heterodimer with high affinity for heregulin growth factors (Sliwkowski et al., 1994). ERBB3 over-expression promotes resistance to therapeutic ERBB2 inhibitors in breast cancer (Garrett et al., 2011). Some proteins commonly annotated as pseudokinases have been found to have phospho-transfer activity. HASPIN, for example, is annotated as a pseudokinase in the ProKinO database because it lacks a DFG motif in the catalytic domain, but it has been shown to phosphorylate histone H3 using a DYT motif instead (Eswaran et al., 2009; Villa et al., 2009). H3 phosphorylation by HASPIN changes chromatin structure and mitotic outcome and is therefore physiologically important (Dai et al., 2005). The existence of biologically active pseudokinases, some of which may have phosphotransferase activity, is one way in which equating kinases with a specific fold, sequence similarity, or enzymatic function is inadequate.

Based on the current work, the most useful definitions of the human kinome are likely to be an expansive 710 domain “extended kinome” that broadly encompasses related sets of folds, sequences and biological functions and a 557 domain “curated kinome” that is most similar in spirit to the original definition by Manning (Manning et al., 2002) nearly two decades ago. The extended kinome is useful for machine learning, chemoproteomics, small molecule profiling and genomic studies in which an expansive view of kinase biology is advantageous. The curated kinome is most useful in the study of kinases as a family of genes with related biochemistry, folds, and cellular function.

### The challenge of unknown knowns

The current study makes clear the magnitude of the problem presented by the “unknown knowns” in kinase biology and biomedicine more generally. In the current context, an unknown known is a relevant piece of experimental data about one or more kinases described in the peer-reviewed literature that cannot be found by querying existing database or a via a standard text-based search on PubMed. A common reason for the inability of human curators and NLP software to find relevant published information is the widespread use of informal and non-uniform identifiers and names, particularly in biochemical studies. Four example, the four members of the p38 kinase family (the α, β, γ, and δ isoforms in the biochemical literature and <u>MAPK14</u>, <u>MAPK11</u>, <u>MAPK12</u> and <u>MAPK13</u> respectively in the HUGO namespace) have in aggregate at least 25 aliases (PRKM11, SAPK2, SAPK2B, p38-2, CSBP1, CSBP2, CSPB1, EXIP, Mxi2, PRKM14, PRKM15, RK, SAPK2A, ERK3, ERK5, ERK-6. ERK6, PRKM12, SAPK-3, SAPK3, MGC99536, PRKM13. SAPK4) often with dashes in different places and various misspellings. In principle, NLP systems should be able to overcome these inconsistencies via robust grounding algorithms, but we find that misspellings, errors in residue numbering, use of mouse names for human proteins and vice versa, and use of ambiguous acronyms remain a substantial barrier to assembly of systematic knowledge about kinases (Bachman et al., 2019; Steppi et al., 2020) and presumably other classes of proteins as well. Moreover, in the scientific literature, findings about kinases are often described at the level of protein families (e.g., MEK, AKT, ERK) or complexes (e.g., mTORC1, PI3K) rather than one or more of their specific protein members (Bachman et al., 2018). For instance, whereas “p38 kinase” is found in ∼42,000 PubMed publications (as of January, 2021), the HUGO standardized names MAPK11-14 are found in only ∼1,500 publications in total. INDRA uses tools such as such a FamPlex (Bachman et al., 2018) to distinguish between knowledge about protein families, complexes and individual proteins in a principled manner. The INDRA networks we have constructed for understudied kinases exploit state-of-the art grounding and FamPlex but much work remains to be done to fully “rediscover” unknown knowns.

The issue of accurate entity grounding is not simply a matter of semantics. Reagents used to measure ERK kinases such as antibodies do not necessarily distinguish between MAPK1/ERK1 and MAPK3/ERK2, and the same is true of antibodies against p38 kinases, particularly phosphospecific antibodies. In the case of ERK1/2 this ambiguity might not be critical, because the two enzymes are generally regarded as functionally redundant (Buscà et al., 2015, 2016). However, the four isoforms of p38 kinases are distinct in patterns of expression, biological function and binding to small molecule inhibitors (Roux and Blenis, 2004). Subject matter experts can sometimes disambiguate these references based on the experimental system, the methods used, or knowledge of past publications on similar topics, but this information is difficult and time consuming for humans to curate and machine reading is not yet sufficiently sophisticated. Automated discovery is also substantially hindered by the inability to read the full text of most copyrighted literature, even if personal and institutional subscriptions are in place. The net result is that much of the information on the kinome that has been collected through careful experimentation and paid for with public funds is not easily available in an organized way to most of the scientific community. We continue to receive feedback about information missing from the web resources described here and make our own serendipitous discoveries in the literature (we welcome help in adding any information we have missed via email to the corresponding author). We speculate that addressing the issue of unknown knowns will have a bigger impact on our understanding the kinome and other multi-gene families than any near-term series of experimental studies; the development of better NLP, curation and knowledge assembly tools should therefore be a high priority.

### The understudied kinome

Following the lead of the NIH IDG consortium, we have defined the understudied kinome as the least-studied one-third of all kinase domains even though the distribution is unimodal. These understudied domains have ∼13-fold fewer INDRA statements on average than well-studied kinases and they are less likely to have small molecule ligands. Among the 182 kinases in the curated kinome that we currently judge to be understudied, 53 have high resolution structures and for 101 commercial activity and/or binding assays exist: CLK3 and WNK3, for example, both have crystal structures and are available as recombinant protein on Reaction Biology and DiscoverX kinase panel. In general, we find that the number of INDRA statements for a kinase correlates well with the number of selective or clinical grade small molecules but that some relatively well-studied kinases do not have high affinity chemical ligands.

We find that at least 175 of the understudied kinases as defined above, are expressed in CCLE cell-lines (the largest cell-line panel analyzed to date) when measured by protein or by mRNA expression (and in many cases by both). We also find that half of all understudied kinases are essential in two or more of the 625 cell-lines annotated in the DepMap (Tsherniak et al., 2017) and 10 are essential in at least two-thirds of DepMap lines. In addition, 27 kinases are among the top ten most mutated kinases in one or more cancer types annotated in TCGA and several others are differentially expressed in disease databases for Alzheimer’s Disease or COPD. These largely indirect findings suggest that a substantial subset of understudied kinases are functional in normal physiology and in disease. This information is of immediate use in studying protein phosphorylation networks and it sets the stage for use of genetic and chemical tools to investigate the functions of specific understudied kinases. Based on available evidence, the possibility exists that some understudied kinases may be valuable as therapeutic targets.

## Supporting information

metadata file

## ACKNOWLEDGEMENTS

This work was funded by grant U24-DK116204 and U01-CA239106 as part of the National Institutes of Health Illuminating the Function of the Understudied Druggable Genome Program. The development of INDRA was funded by DARPA grants W911NF-15-1-0544 and W911NF018-1-0124. We thank other members of the Understudied Kinase Consortium including PIs Gary Johnson (UNC), Tim Wilson (UNC), Ben Major (WUSL) and Reid Townsend (WUSL) as well Cat Luria (HMS), Mike East (UNC), Tudor Oprea (UNM) and Jeremy Muhlich (HMS); we thank Tony Hunter (Salk Institute) for reviewing the updated protein kinase list and Yuan Wang and Jeremy Jenkins (Novartis) for providing the Bayesian Affinity Fingerprint vectors. The results on Azheimer’s disease published here are in part based on data obtained from the AMP-AD Knowledge Portal (https://adknowledgeportal.synapse.org/). Study data were provided by the Rush Alzheimer’s Disease Center, Rush University Medical Center, Chicago. Data collection was supported through funding by NIA grants P30AG10161, R01AG15819, R01AG17917, R01AG30146, R01AG36836, U01AG32984, U01AG46152, the Illinois Department of Public Health, and the Translational Genomics Research Institute. Additional data were generated from postmortem brain tissue collected through the Mount Sinai VA Medical Center Brain Bank and were provided by Dr. Eric Schadt from Mount Sinai School of Medicine.

## OUTSIDE INTERESTS

PKS is a member of the SAB or Board of Directors of Applied Biomath and RareCyte Inc and has equity in these companies; he is a member of the SAB of NanoString Inc.. In the last five years the Sorger lab has received research funding from Novartis and Merck. BMG and JAB have received consulting fees from TwoSix Labs, LLC. Sorger, Gyori and Bachman declare that none of these relationships are directly or indirectly related to the content of this manuscript. Other authors declare that they have no outside interests.

## METHODS

### Classification of the “extended kinome” and defining the “curated kinome”

To obtain a list of kinases from UniProt all human proteins annotated to have kinase activity were extracted and filtered based on (i) interaction with ADP/ATP; (ii) presence of a kinase domain; 3) membership in a kinase family (lists of kinase domains and kinase families are available in supplementary material). To identify human kinase sequences that belong to the Protein Kinase Like (PKL) fold, 710 sequences annotated as “kinase” in UniProt were first subjected to a similarity search against well curated ePK profiles to identify and separate out the 8 canonical ePK groups (Eswaran et al., 2009; Manning et al., 2002; Talevich et al., 2011). eLKs were identified based on detectable sequence similarity with one or more of the ePK sequences. Sequences that share no detectable sequence similarity to ePKs were classified as aPKs. For predicted aPKs, crystal structures of the protein itself or of the closest homolog were inspected manually to check if the kinase domain adopts a canonical ePK fold. Additional support for this classification was obtained by calculating a Hidden Markov Model (HMM)-based distance score between the Pfam domains (Huo et al., 2017) and the presence/absence of key structural features distinguishing ePKs, eLKs and aPKs, as described previously(Kannan and Neuwald, 2005; Kannan et al., 2007). Kinases with a kinase domain distinct in sequence and fold from the known protein kinases were classified as “Unrelated to Protein Kinase” (uPK). A subset of sequences that satisfied none of the above criteria i.e., no detectable sequence similarity to ePKs, no clear kinase function and no homologous crystal structures, were grouped into the *unknown protein kinase* category (Unknown). All kinases annotated to have a PKL fold were included in the curated kinome. STK19 was also included in the curated kinome despite its unknown fold since it is known to be serine/threonine kinase active against peptide substrates (Yin et al., 2019).

### Curation of INDRA statements and generation of INDRA networks

INDRA uses natural language processing (NLP) systems to extract mechanistic information from literature as well as databases and represents them in a standardized format as previously described (Gyori et al., 2017). The corpus of statements about kinases was assembled by processing all accessible MEDLINE abstracts, as well as the PubMed Central Open Access and Author’s Manuscript subsets using multiple machine-reading systems integrated with INDRA. Statements originating from machine reading were combined with ones extracted from structured knowledge bases including Pathway Commons and SIGNOR. To count the number of corresponding INDRA Statements, and construct kinase-specific networks, we queried this knowledge based for each kinase (identified by its HGNC identifier) using the script “get_kinase_interactions.py”. We generated kinase-specific networks using the INDRA CX assembler (indra.assemblers.cx module) and uploading each network to NDEx (ndexbio.org) where they can be browsed and queried. We applied a “semantic hub” layout to these networks which positions the biological entities around a kinase of interest based on their type (e.g., protein, small molecule, biological process), and role as upstream or downstream interactor. The python scripts to collect INDRA statements, calculate kinase-specific statement counts, and assemble networks are available on the Github repository (https://github.com/IDG-Kinase/indra_analysis).

### Small molecule selectivity calculations

The specificity of small molecules was calculated according to the *selectivity score* (Moret et al., 2019), which uses multiple parameters to assess selectivity: (i) the absolute affinity for the “on” target; ii) the differential affinity between the “on” and “off” targets of each kinase; (iii) the p-value of the difference between the distributions of “on” and “off” targets; (iv) the research bias – a score indicating how broadly a compound has been tested for off-targets. The selectivity score was divided in four tiers; Most Selective (MS), Semi Selective (SS), Polyselective (PS) and Unknown (UN). MS levels are defined as an absolute affinity of Kd <100 nM (at least two measurements); a differential affinity of 100 (i.e., the affinity of the compound for the “on” target is 100 times greater than for the “off” targets), a p-value ≤ 0.1 and a research bias <0.2; SS levels are defined as an absolute affinity of Kd<1 µM (at least 4 measurements), a differential affinity of 10, a p-value ≤0.1 and research bias <0.2; PS levels are defined as an absolute affinity Kd< 9000 nM, differential affinity of 1 (e.g., equal affinity for “on” and “off” targets) and research bias <0.2; UN levels are defined as an absolute affinity Kd< 9000 nM and differential affinity of 1.

### CCLE analysis

The data RNA dataset “CCLE_RNAseq_genes_rpkm_20180929.gct.gz” was downloaded from the CCLE portal (https://portals.broadinstitute.org/ccle/data) and analyzed with the script “analyzing_CCLE_data.r”. The maximum expression value over all cell-lines was calculated and plotted (Figure 3A). Genes were considered “expressed” if the maximum RPKM was ≥1. The mass spectrometry dataset “protein_quant_current_normalized.csv” was downloaded from the DepMap portal (https://depmap.org/portal/download/) and analyzed with the script “analyzing_CCLE_data.r”. Proteins for which one or more peptides were detected in this dataset were considered to be expressed.

### Determination of Essential Kinases through the Dependency Map

The preprocessed results of genome-wide CRISPR knockout screens were obtained from the DepMap 19Q4 Public data release (https://depmap.org/portal/download/). The results of the screens were processed as described by Dempster et al (Dempster et al., 2019). For each kinase, cell-lines with a CERES score >0.5 were classified as dependent and the number of dependent cell-lines for each kinase was then tallied.

### TCGA analysis

TCGA PanCan gene expression and mutation frequency data was obtained from cBioPortal (Cerami et al., 2012; Gao et al., 2013). To identify kinases with abnormal expression in tumors, tumor types with at least 10 paired normal tissue samples were analyzed. For each kinase, the fold change of its median expression in either all tumor tissues (general PanCan analysis) or the individual tumor tissue over its median expression in the paired healthy tissues was calculated. P-value from Wilcoxon-Mann-Whitney two-sided test was calculated based on the distributions of gene expression in tumor and healthy tissues in each tumor type. Adjusted p-values were calculated using the Benjamini-Hochberg procedure. To identify kinases heavily mutated in cancer, the number of patient samples with mutation or gene fusion was counted and normalized to the total number of patient samples (10953 samples).

### AMP-AD analysis

Preprocessed count matrices of AMP-AD consortium RNA-seq data were downloaded from the AMP-AD Synapse directory (https://adknowledgeportal.synapse.org/). In summary, these counts were derived from raw reads using the STAR aligner(Dobin et al., 2013) and the Gencode v24 human genome annotation. In our analysis, we included all Alzheimer’s disease (AD) patients from the Mount Sinai VA Medical Center Brain Bank (MSBB) and the Religious Orders Study and Memory and Aging Project (ROSMAP) study(Mostafavi et al., 2018) for which RNA-seq data from post-mortem brain was available and their age at death and Braak stage were known. Differential expression analysis was performed using the R package DESeq2(Love et al., 2014). We fitted a generalized linear model to the expression of each gene using the Braak stage as independent variable and adjusted for age at death and study batch effect by including them as covariates. We used the Wald test implemented in DESeq2 to extract differentially expressed genes between early (Braak stages 1 and 2) and late (5 and 6) AD cases. Effect sizes were moderated using the R package apeglm (Zhu et al., 2019).

### COPD differential expression analysis

Preprocessed dataset combining 5 datasets from GEO and 2 from ArrayExpress was downloaded from https://figshare.com/articles/Meta-analysis_of_Gene_Expression_Microarray_Datasets_in_Chronic_Obstructive_Pulmonary_Disease/8233175. Data was preprocessed as described in Rogers et al (Rogers et al., 2019). Raw expression data was processed by generalized least squares (GLS) weighted models to account for heterogeneity between datasets. A Likelihood Ratio Test was used to identify differentially expressed genes. Genes with significant (adjusted p-value <0.5) differential expression in COPD versus healthy individuals and that are within the two-tailed 10% and 90% quantile were identified as genes of interest. Relative expression of these differentially expressed genes was calculated as the effect size of the GLS estimates of the individuals with COPD and healthy individuals.

### Library analysis

Library OKL2.0 was analyzed as described previously (Moret et al., 2019). RDKit (www.rdkit.org) was used for fingerprinting and chemical similarity calculations.

## SUPPLEMENTARY FIGURE LEGENDS

**Figure S1 related to** Figure 1 **- Kinase domains in the standard list**

**(A)** Pie chart of kinases in the Manning and KinHub lists divided into kinase groups as conventionally defined. Letter coding explanation can be found of main figure legend 1C. **(B)** Number of compounds that target understudied kinases as determined in a recent large-scale chemoproteomic assay (Klaeger et al., 2017).

**Figure S2 related to** Figure 3 **- INDRA network for WEE2**

**(A)** A partial network (upper panel) and statement table (lower panel) generated by INDRA for the understudied kinase WEE2. Table contains full quotes from literature. **(B)** Comparison of number of INDRA statements (X-axis) and TIN-X novelty score (Cannon et al., 2017) (Y-axis) for all domains in the curated human kinome. The number of INDRA statements correlates with TIN-X novelty score with a Pearson’s correlation coefficient of r = 0.81. The bottom third of domains having the least knowledge according to both INDRA and TIN-X are highlighted in pink and constitute the understudied kinome as defined in this manuscript. In **panel B** the Pharos target designation (solid colors) and IDG status (shape) are shown; in **panel C**, the fill color represents the maximum selectivity of a small molecule compound known to bind to each kinase. See text for details.

**Figure S3 related to** Figure 6 **- Differential mRNA expression of kinases in selected TCGA datasets**

**(A)** Differential mRNA expression for both the understudied and the well-studied kinases in colon adenocarcinoma (COAD) based on data in TCGA and accessed via c-BioPortal.82. HGNC symbols and cancer type abbreviations for selected outlier genes. **(B)** Depicted is the alteration frequency in lymphoid neoplasm diffuse large B-cell lymphoma (DLBCL) for understudied kinases (darks bars) and well-studied kinases (light bars). Both fusion (magenta) and mutations (black/grey) are indicated.

**Figure S4 related to** Figure 8 **– clinical compounds potentially binding dark kinases**

Kinase inhibitors in clinical development and FDA-approved small molecule therapeutics targeting understudied kinases for which binding is scored as *most selective (*MS) or *semi selective (*SS) based on data from the Small Molecule Suite (SMS). Seventeen understudied kinases are targeted in total according to SMS. Experimental validations of the compounds binding these targets were performed and twenty interactions were confirmed. Single asterisk: interaction did not bind in our assay. Double asterisk: no commercial assay available at this time.

**Figure S5 related to** Figure 8 **– Clustering of kinases by binding pocket based on BAFP and mapped to the Coral dendrogram**

**(A)** BAFP clusters visualized on the Coral kinase dendrogram. **(B)** The nine labelled BAFP clusters (denoted by color and labelled 1-9) projected on one Coral kinase dendrogram. Understudied kinases in each cluster are colored black.

**Figure S6 related to** Figure 8 **-Kinase domains with high resolution structures**

Number of structures of the kinase domain in Protein Data Bank (PDB) for both well-studied kinases (orange) and understudied kinases (purple) sorted in descending order. Top understudied kinases with high number of kinase domain structures are labeled.

**Figure S7 related to** Figure 9 **– Performance of the commercial Optimal Kinase Library.**

**(A)** Maximum selectivity of compounds in the commercial Optimal Kinase Library for the curated kinome. **(B)** Maximum selectivity of compounds in the theoretical Optimal Kinase Library for the curated kinome. **(C)** Distribution of the Target Affinity Spectrum (TAS) similarity. **(D)** Comparison of structural similarity and TAS similarity. **(E)** Distribution of structural selectivities in the Optimal Kinase Library. **(F)** comparison for the phenotypic fingerprint (PFP) correlation and the number of assays that the compound pairs had in common

## FOOTNOTES FOR SUPPLEMENTARY TABLES

**Supplementary Table 1 – The extended kinome**

This table describes available information about the 710 kinase domains in the extended kinome. Each domain is annotated with the following pieces of information: gene_id (NCBI gene ID); UniProtEntry (Uniprot ID, Uniprot Entry name and domain index if the kinase contains multiple kinase domains) Entry (the unique and stable short-form Uniprot ID as a number); Entry name (Mnemonic identifier to UniprotKB entry); Gene names (names of genes encoding this protein as obtained from Uniprot), Protein names (full name of the protein provided by UniProt), HGNC ID, HGNC_name (the official gene symbol approved by HGNC), Approved name (the full gene name approved by HGNC), IDG_dark (value of 0 or 1 denoting whether part of the IDG dark kinase list), KinHub (value of 0 or 1 denoting whether the domain is on the KinHub list), Manning (value of 0 or 1 denoting whether domain is on the Manning list), Group (membership to one of the ten kinase groups), Family (membership in the kinase families), Uniprot_kinaseactivity (value of 0 or 1 denoting whether domain is on the curated UniProt kinase list), PfamDomain, DomainStart (first residue number of the kinase domain according to UniProt), DomainEnd (last residue number of the kinase domain according to UniProt), ProKinO (value of 0 or 1 denoting wehther the domain is in ProKinO), New_Annotation (further classification of the protein fold as ePK, eLK, Atypical, Unrelated to Protein Kinase or Unknown), Fold (primary classification of the protein fold: protein kinase like – PKL -, unrelated, UPK or unknown), Pseudokinase? (Yes or No annotation to whether the kinase is a pseudokinase according to ProKinO), Annotation_Score (number to reflect the amount of aggregated information from multiple databases), INDRA_network (URL of the interactive INDRA network on NDEx).

**Supplementary Table 2 – The curated kinome**

A table describing data about the 556 kinase domains in the curated kinome with a PKL fold (plus STK19). Each domain is annotated with all information in Supplementary Table 1 about its identifiers (NCBI gene_id, HGNC identifiers, and UniProt identifiers), inclusion and exclusion criteria based on different kinase lists (Manning, KinHub, kinase group and kinase family according to KinHub, curated UniProt kinase list, IDG dark kinases, ProKinO, pseudokinase), protein fold, and the URL for its INDRA network on NDEx. Each kinase domain also has the following additional annoations: **(i)** amount of existing information (n_indra_statement: number of INDRA statements; TIN-X_Score; tdl (target development level from Pharos); **(ii)** whether the kinase is understudied (stat_dark_num: value of 0 or 1 denoting whether a kinase is understudied based on number of INDRA statement and TIN-X_Score); **(iii)** PDB structures (PDBID: PDB IDs for any structures of the kinase domain; num_pdb: the total number of pdb structures of the kinase domain); **(iv)** number of MS/SS compounds (num_MSSS_cmpd); **(v)** availability of commercial activity assays (rb_name: the name of the kinase on Reaction Biology (http://www.reactionbiology.com); rb_variants: the phosphorylated form or protein complex available for assay on Reaction Biology; kinomescan_name: the name of the kinase on DiscoverX (https://www.discoverx.com/home); kinomescan_variants: the phosphorylated form or protein complex available for assay on the DiscoverX kinase panel; commercial_assay: value of 0 or 1 denoting whether a Reaction Biology or KinomeScan assay is available); **(vi)** biological relevance and disease implications (num_dep: number of dependent cell-lines on DepMap; AMPAD: value of 0 or 1 denoting whether the kinase is differentially expressed in Alzheimer patients; TCGA: value of 0 or 1 denoting whether the kinase is differentially expressed in any cancer type, among the top 10 most frequently mutated kinases in any cancer type, or among the top 20 most frequently mutated kinases of all cancers; COPD: value of 0 or 1 denoting whether the kinase is differentially expressed in COPD patients). The metadata for this table can be found in ‘meta_data.xlxs’

**Supplementary Table 3 – Clinical compounds targeting understudied kinases**

A table with the affinity values of compounds in clinical development (phase 1-3) or approved drugs that have been shown to target understudied kinases based on available data in Small Molecule Suite (http://smallmoleculesuite.org). The compounds are annotated with IC50_Q1 (the affinity value per understudied kinase), HGNC_symbol (official HGNC symbol of understudied kinase), compound_max_phase (the latest stage of clinical development), compound_first_approval (year of first approval if compound is an approved therapeutic), compound_chembl_id (ChEMBL identifier of compound).

