## Supplementary Figures for "A resource for exploring the understudied human kinome for research and therapeutic opportunities"

#### FIGURE S1

A

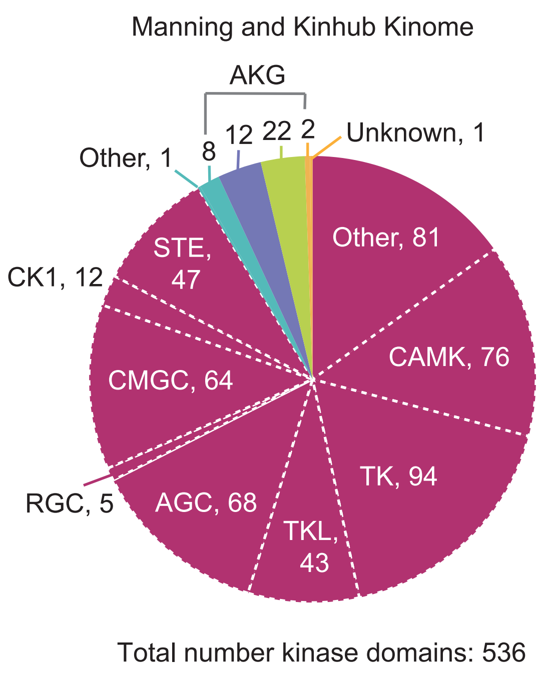

Manning and Kinhub Kinome
AKG
Other, 1
8
12
22
2
Unknown, 1
Other, 81
STE, 47
CK1, 12
CAMK, 76
CMGC, 64
TK, 94
RGC, 5
AGC, 68
TKL, 43
Total number kinase domains: 536


B

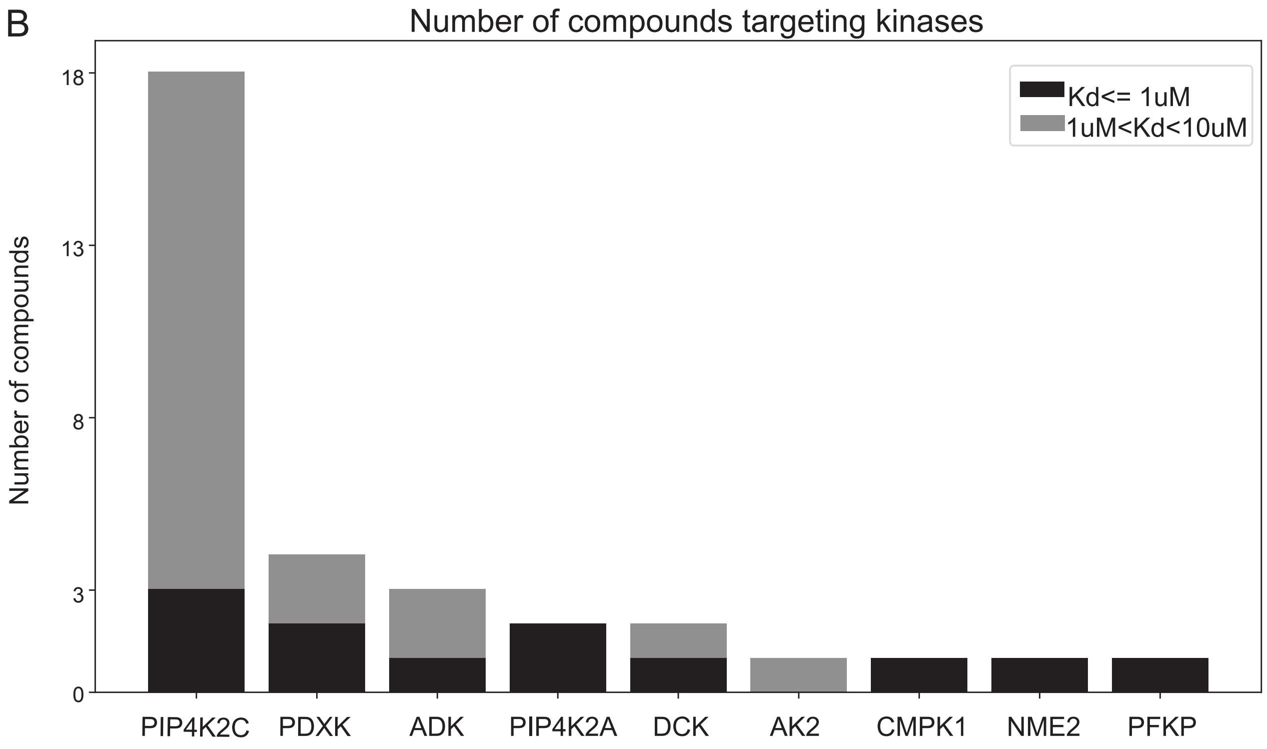

Number of compounds targeting kinases
Number of compounds
18
13
8
3
0
Kd<= 1uM
1uM<Kd<10uM
PIP4K2C
PDXK
ADK
PIP4K2A
DCK
AK2
CMPK1
NME2
PFKP

#### FIGURE S2

**a.** WEE2 INDRA network

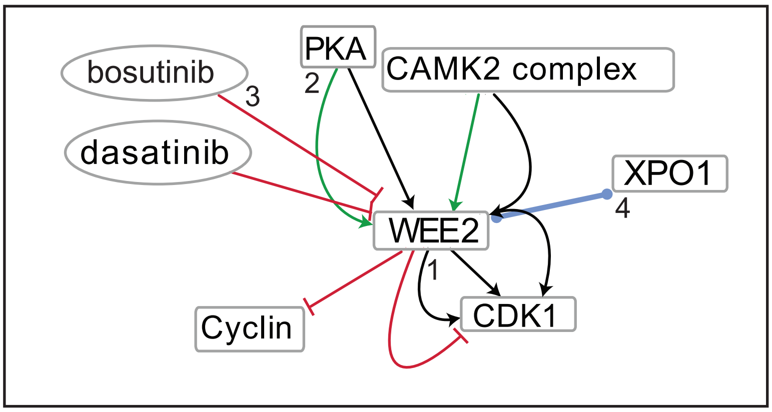


| Edge index | Connection Type | Statement | Evidence | Evidence Type | Link |
|---|---|---|---|---|---|
| 1 | Modification | Phosphorylation(WEE2(), CDK1()) | CaMKII regulates resumption of the cell cycle through phosphorylation of Wee1B, which in turn phosphorylates and inhibits Cdc2 | literature | PMID: 23287037 PMID: 21454751 PMID: 19550110 |
| 2 | Activation | Activation(PKA(), WEE2()) | Furthermore, PKA phosphorylates and activates the CDK1 inhibitor Wee1B which is the oocyte-specific Wee1 isoform. | literature | PMID: 23805152 |
| 3 | Inhibition | Inhibition(Bosutinib(), WEE2()) | - | database | Small Molecule Suite |
| 4 | Complex | Complex(XPO1(), WEE2()) | Together with the localization data, these findings document that Wee1B binds to Crm1 through the identified NES sequence and point mutations in this region disrupt this interaction | literature | PMID: 20083600 |
| 5 | … | … | … | … | … |

**b.** Knowledge about the kinome extracted by INDRA and TIN-X

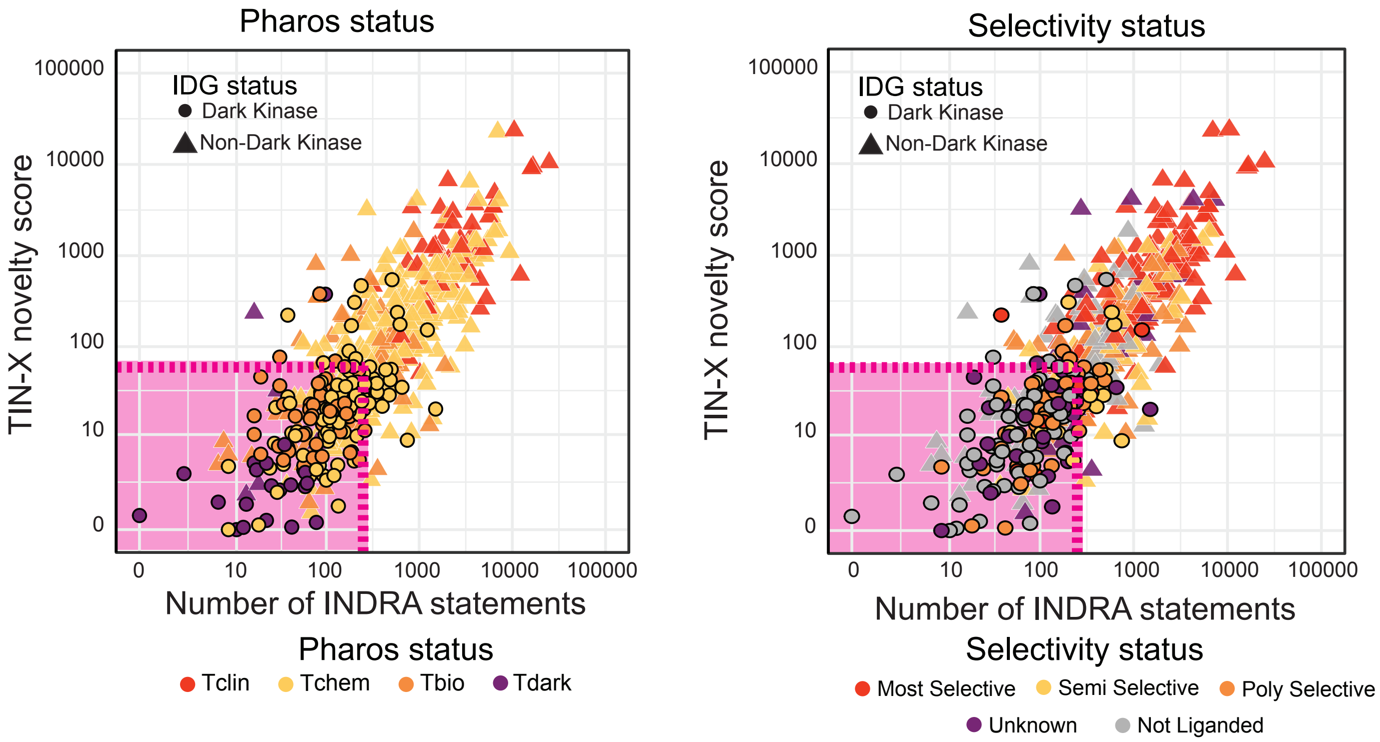

#### FIGURE S3

A

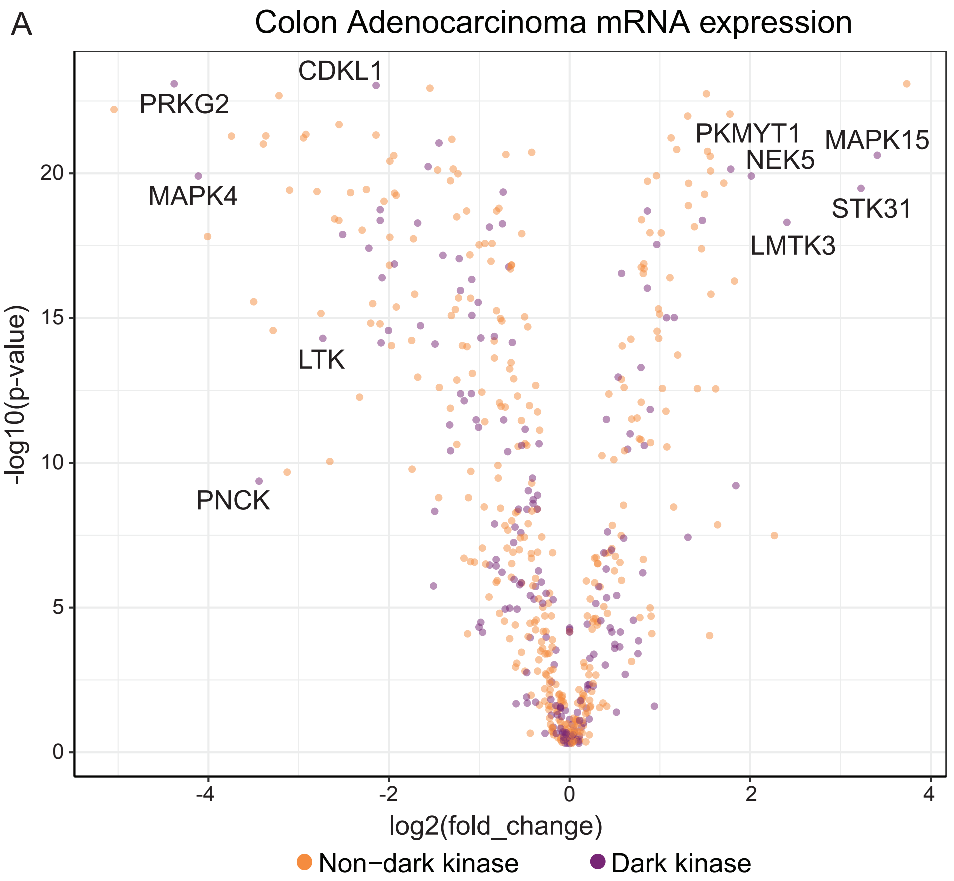
Colon Adenocarcinoma mRNA expression
CDKL1
PRKG2
MAPK4
LTK
PNCK
PKMYT1
MAPK15
NEK5
STK31
LMTK3
-log10(p-value)
0
5
10
15
20
-4
-2
0
2
4
log2(fold_change)
Non−dark kinase
Dark kinase

B

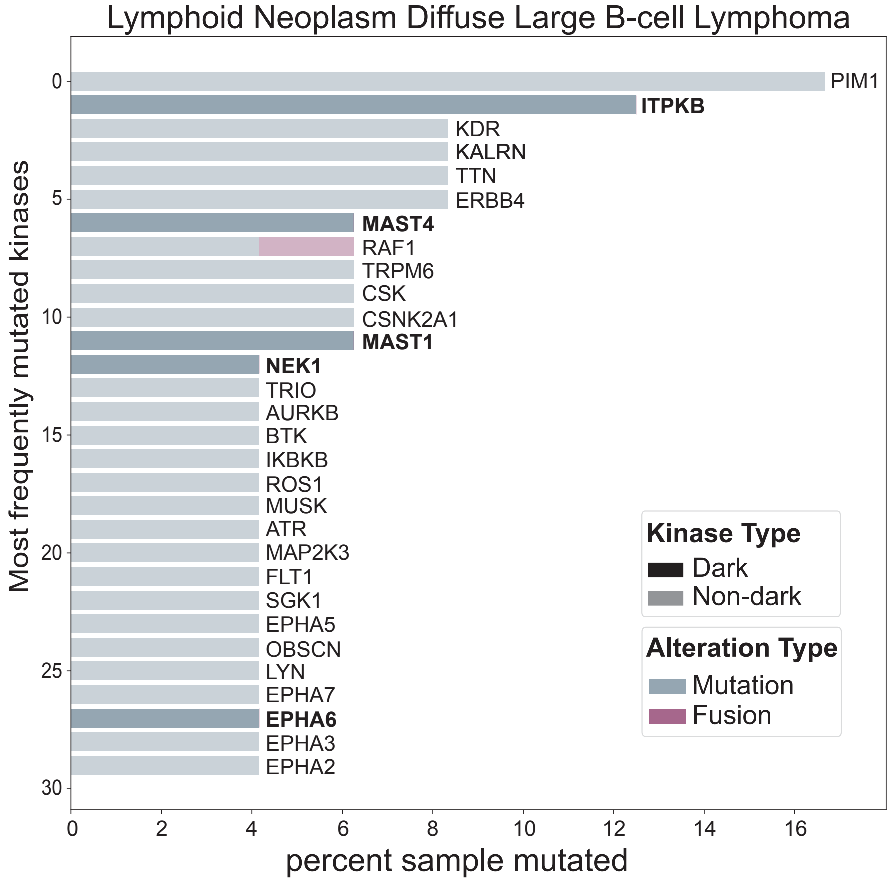
Lymphoid Neoplasm Diffuse Large B-cell Lymphoma
PIM1
ITPKB
KDR
KALRN
TTN
ERBB4
MAST4
RAF1
TRPM6
CSK
CSNK2A1
MAST1
NEK1
TRIO
AURKB
BTK
IKBKB
ROS1
MUSK
ATR
MAP2K3
FLT1
SGK1
EPHA5
OBSCN
LYN
EPHA7
EPHA6
EPHA3
EPHA2
Most frequently mutated kinases
0
5
10
15
20
25
30
0
2
4
6
8
10
12
14
16
percent sample mutated
Kinase Type
Dark
Non-dark
Alteration Type
Mutation
Fusion

#### FIGURE S4

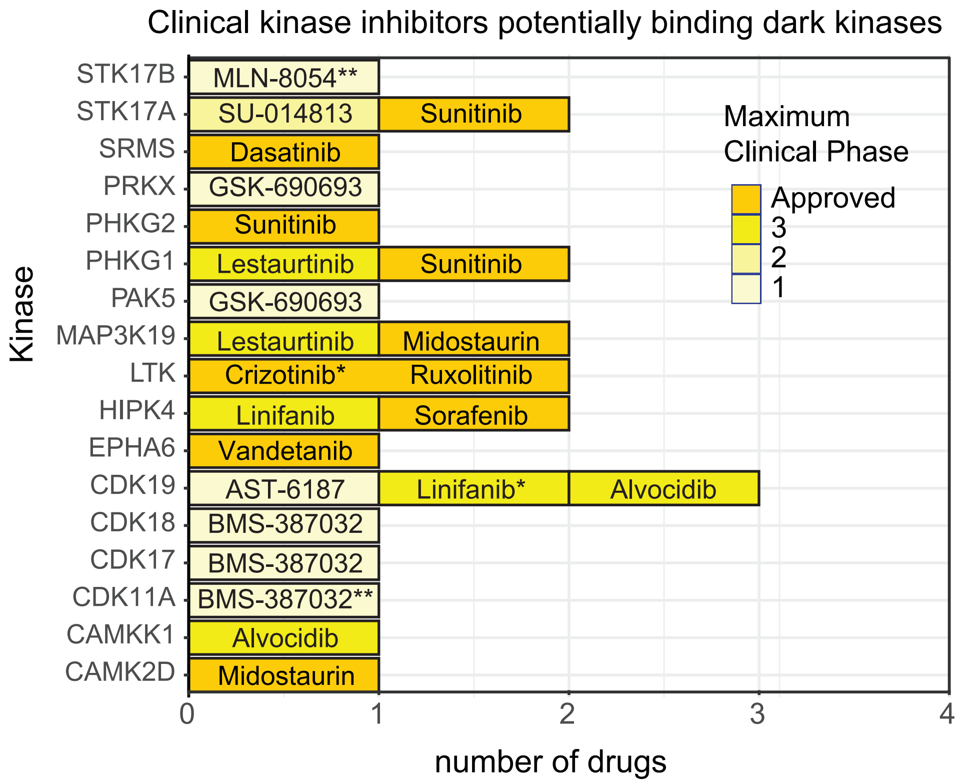

Clinical kinase inhibitors potentially binding dark kinases
Kinase
STK17B
STK17A
SRMS
PRKX
PHKG2
PHKG1
PAK5
MAP3K19
LTK
HIPK4
EPHA6
CDK19
CDK18
CDK17
CDK11A
CAMKK1
CAMK2D
MLN-8054**
SU-014813
Sunitinib
Dasatinib
GSK-690693
Sunitinib
Lestaurtinib
Sunitinib
GSK-690693
Lestaurtinib
Midostaurin
Crizotinib*
Ruxolitinib
Linifanib
Sorafenib
Vandetanib
AST-6187
Linifanib*
Alvocidib
BMS-387032
BMS-387032
BMS-387032**
Alvocidib
Midostaurin
Maximum Clinical Phase
Approved
3
2
1
0
1
2
3
4
number of drugs

### FIGURE S5

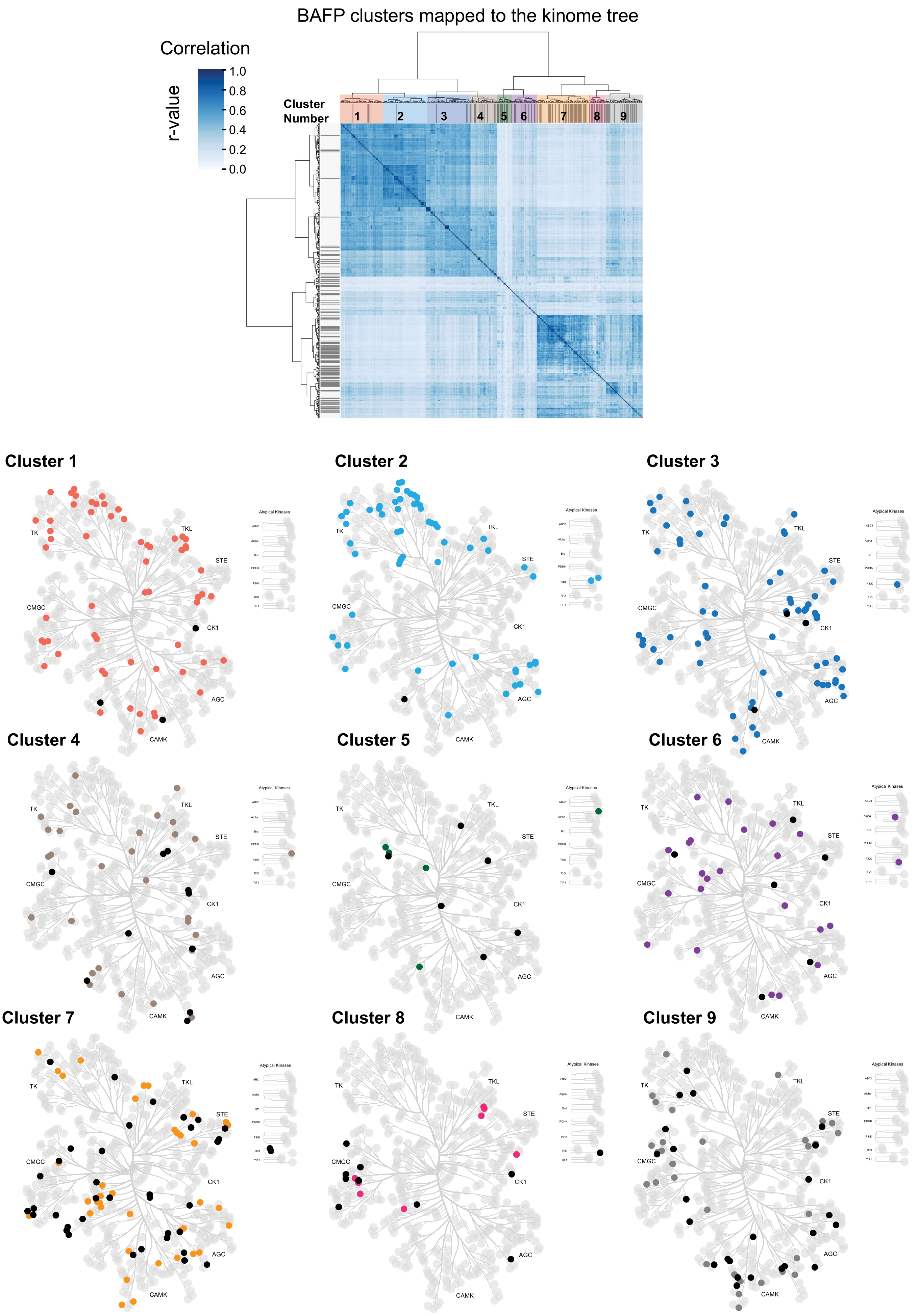
BAFP clusters mapped to the kinome tree
Correlation
r-value
1.0
0.8
0.6
0.4
0.2
0.0
Cluster Number
1
2
3
4
5
6
7
8
9
Cluster 1
Cluster 2
Cluster 3
Cluster 4
Cluster 5
Cluster 6
Cluster 7
Cluster 8
Cluster 9
TK
TKL
STE
CMGC
CK1
AGC
CAMK
Atypical Kinases

#### FIGURE S6

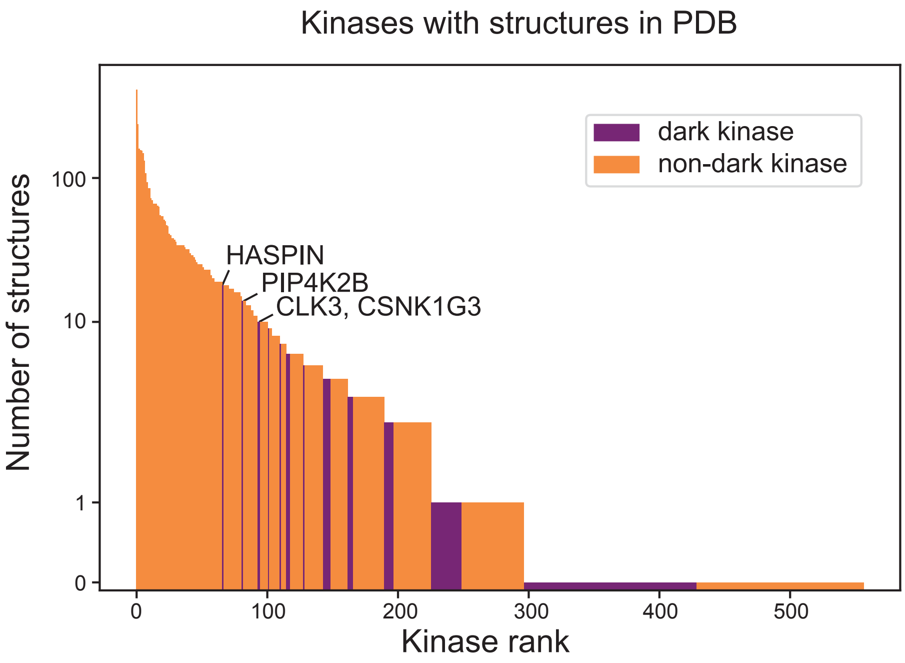
Kinases with structures in PDB
dark kinase
non-dark kinase
HASPIN
PIP4K2B
CLK3, CSNK1G3
Number of structures
100
10
1
0
0
100
200
300
400
500
Kinase rank

### FIGURE S7

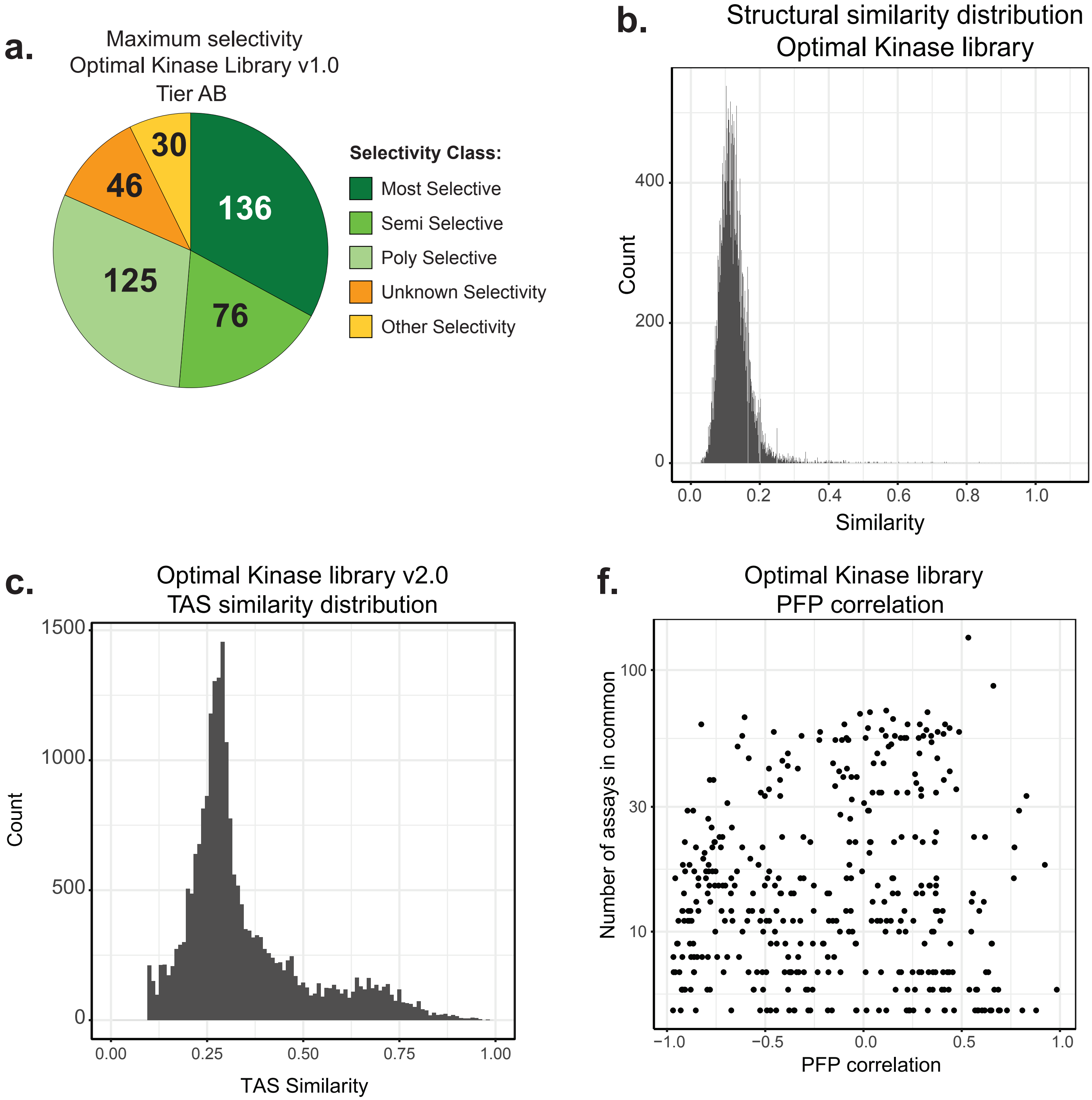
a.
Maximum selectivity
Optimal Kinase Library v1.0
Tier AB
30
46
136
125
76
Selectivity Class:
Most Selective
Semi Selective
Poly Selective
Unknown Selectivity
Other Selectivity
b.
Structural similarity distribution
Optimal Kinase library
Count
400
200
0
0.0
0.2
0.4
0.6
0.8
1.0
Similarity
c.
Optimal Kinase library v2.0
TAS similarity distribution
Count
1500
1000
500
0
0.00
0.25
0.50
0.75
1.00
TAS Similarity
f.
Optimal Kinase library
PFP correlation
Number of assays in common
100
30
10
−1.0
−0.5
0.0
0.5
1.0
PFP correlation
